# A Mathematical Model of Glomerular Fibrosis in Diabetic Kidney Disease to Predict Therapeutic Efficacy

**DOI:** 10.1101/2023.04.02.535270

**Authors:** Haryana Y. Thomas, Ashlee N. Ford Versypt

## Abstract

1

**Background:** Glomerular fibrosis is a tissue damage that occurs within the kidneys of chronic and diabetic kidney disease patients. Effective treatments are lacking, and the mechanism of glomerular damage reversal is poorly understood.

**Method:** A mathematical model suitable for hypothesis-driven systems pharmacology of glomerular fibrosis in diabetes was developed from a previous model of interstitial fibrosis. The adapted model consists of a system of ordinary differential equations that models the complex disease etiology and progression of glomerular fibrosis in diabetes.

**Results:** Within the scope of the mechanism incorporated, advanced glycation end products (AGE)—matrix proteins that are modified due to high blood glucose—were identified as major contributors to the delay in the recovery from glomerular fibrosis after glucose control. The model predicted that inhibition of AGE production is not an effective approach for accelerating the recovery from glomerular fibrosis. Further, the model predicted that treatments breaking down accumulated AGE is the most productive at reversing glomerular fibrosis. The use of the model led to the identification that glucose control and aminoguanidine are ineffective treatments for reversing advanced glomerular fibrosis because they do not remove accumulated AGE. Additionally, using the model, a potential explanation was generated for the lack of efficacy of alagebrium in treating advanced glomerular fibrosis, which is due to the inability of alagebrium to reduce TGF-β.

**Impact:** Using the mathematical model, a mechanistic understanding of disease etiology and complexity of glomerular fibrosis in diabetes was illuminated, and then hypothesis-driven explanations for the lack of efficacy of different pharmacological agents for treating glomerular fibrosis were provided. This understanding can enable the development of more efficacious therapeutics for treating kidney damage in diabetes.

## 2 INTRODUCTION

Diabetic kidney disease (DKD) is a significant global health problem. With over 135 million patients and a yearly incident rate of over 2 million new cases, the burden of DKD is rising (Deng et al., 2021). DKD is a chronic disease for which treatments that can prevent or completely reverse kidney damage are lacking.

DKD is a complex disease that has multiple factors that cause the development and progression of kidney damage. Although hyperglycemia is the initial stimulus for the development of kidney damage in diabetes, the continued progression of damage is mediated by a host of other stimuli, such as advanced glycation end products (AGE), hypertension, angiotensin, and enhanced oxidative stress (Zhang et al., 2021). These stimuli work both dependently and independently through mechanical and chemical signaling to induce damage such as kidney fibrosis, proteinuria, and glomerular filtration rate decline, ultimately resulting in kidney failure (Thomas and Ford Versypt, 2022). Consequently, treatments must be comprehensive to target the multifaceted nature of DKD. Current treatment for DKD consists of approaches to reduce the risk of the development and progression of kidney damage depending on the stage of DKD. If diagnosed very early, lifestyle changes such as nutrition, exercise, and control of glycemic and lipid levels and blood pressure can reduce the risk of DKD progression (Guedes and Pecoits-Filho, 2022). As DKD progresses, inhibitors of the renin-angiotensin-aldosterone system (RAAS) and sodium-glucose cotransporter-2 (SGLT2) are the suggested treatments (Guedes and Pecoits-Filho, 2022). RAAS and SGLT2 inhibitors, although not cures for DKD, can significantly reduce the risk of further progression of DKD. However, once DKD has advanced, the efficacy of known therapeutic approaches becomes limited (Pillai and Fulmali, 2023).

One of the reasons for the limited efficacy of therapeutics for advanced DKD is the concept of glycemic memory, i.e., even after blood glucose has been regulated, kidney damage continues to progress (Testa et al., 2017; Xu and Guo, 2017; Yamazaki et al., 2021). Glycemic memory was observed in patients who received pancreatic transplants to restore regulation of their blood glucose levels (Fioretto et al., 1998). Although blood glucose normalization was achieved relatively quickly, full recovery from kidney damage was only observed ten years after pancreatic transplant (Fioretto et al., 1998). AGE, enhanced oxidative stress, and epigenetic changes have been strongly implicated as independently or interdependently being responsible for glycemic memory (Drzewoski et al., 2009; Testa et al., 2017; Xu and Guo, 2017; Yamazaki et al., 2021). Epigenetic modifications are still in basic research, while AGE and oxidative stress have been hypothesized to form a vicious cycle that is the major contributor to glycemic memory (Xu and Guo, 2017).

Treatments targeting AGE have been developed but have not entered the market as additional therapies for managing DKD. AGE formation inhibitors, such as aminoguanidine and pyridoxamine, had mixed results in clinical trials (Bolton et al., 2004; Williams et al., 2007; Rabbani et al., 2009). Alagebrium, an AGE crosslink breaker, made it to human clinical trials but was not approved as a viable treatment approach (Toprak and Yigitaslan, 2019).

Questions associated with glycemic memory remain unanswered, such as why glucose control takes years to recover from kidney damage, what approaches can accelerate the recovery from kidney damage, and why treatments for AGE —species implicated in glycemic memory—are not efficacious. To provide explanations for these questions and thus shed more light on hypothesized mechanisms governing glycemic memory, we developed a mechanistic mathematical model of diabetes-induced kidney damage, specifically a model of glomerular fibrosis, by adapting a previous model of interstitial fibrosis in lupus (Hao et al., 2014). We used the core pieces of the previous model, gathered data from the literature, and adapted the model to the mechanisms of disease progression relevant to diabetic glomerular fibrosis. The new model was then used to explore explanations for delayed recovery from glomerular fibrosis and lack of therapeutic efficacy and suggest approaches to accelerate recovery.

## 3 METHODS

### 3.1 Mechanism identification

In the interstitial fibrosis model (Hao et al., 2014), fibrosis occurs through the recruitment of macrophages that cause the activation of resident cells, which then produce excess quantities of extracellular matrix (ECM), resulting in fibrosis. To adapt the model, we compared the mechanistic details of the lupus interstitial fibrosis model to published data relevant to diabetic glomerular fibrosis from *in vitro* and *in vivo* experimental studies and well-established protein-enzyme interactions. The literature-supported mechanism for the progression of glomerular fibrosis in diabetes is shown in Figure 1 and described below.

**Figure 1.**
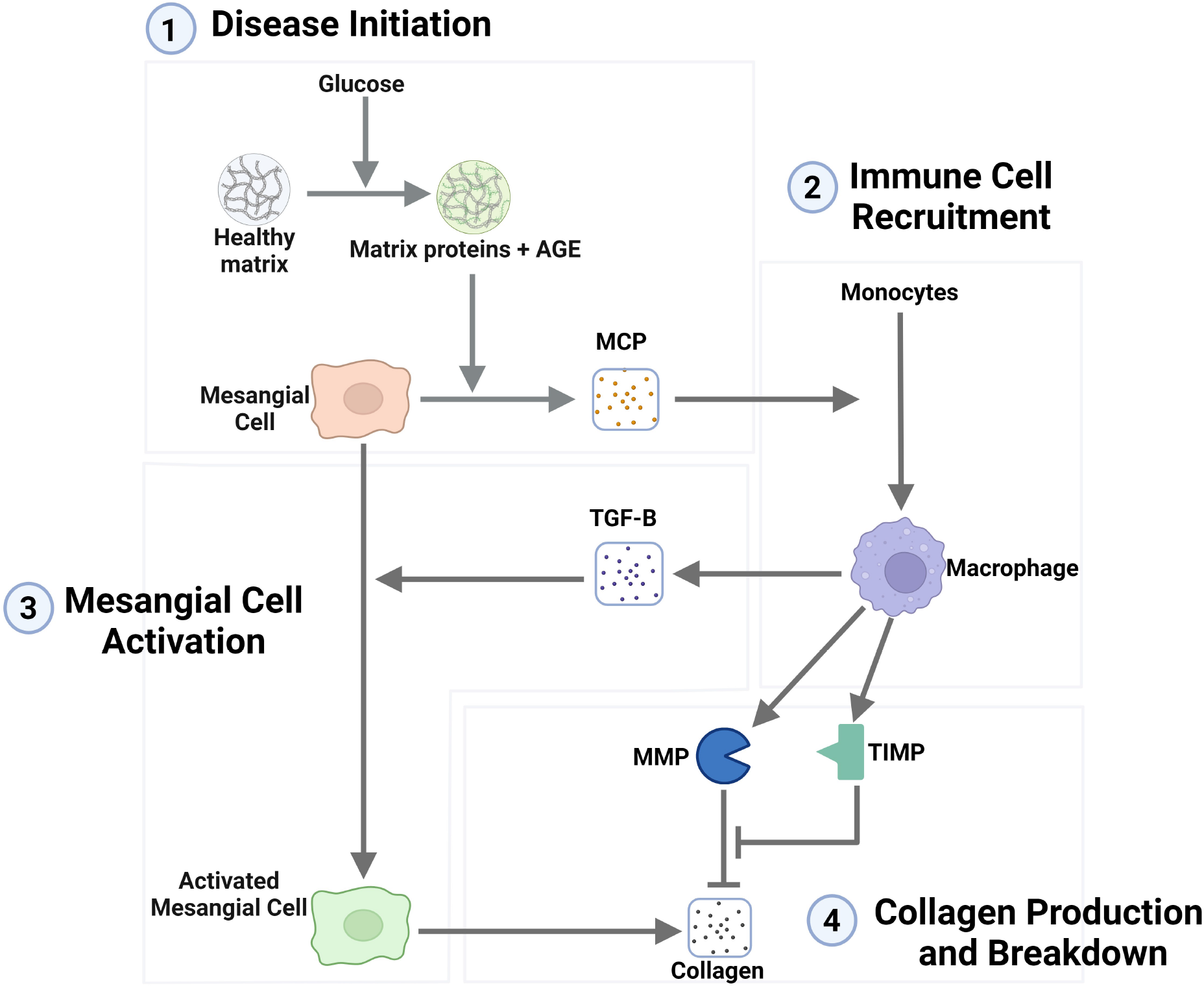
Network representation of the mechanism of the progression of glomerular fibrosis in diabetes consisting of four steps: (1) disease initiation, (2) immune cell recruitment, (3) mesangial cell activation, and (4) collagen production and breakdown. Degradation terms are not shown in this figure. Arrows toward species represent their production, and arrows towards other arrows represent the activation of a process. Flat-ended lines towards other lines represent inhibition, and flat-ended lines towards species represent degradation. Abbreviations: AGE, advanced glycation end products; MCP, monocyte chemoattractant protein; TGF-β, transforming growth factor–β; MMP, matrix metalloproteinase; TIMP, tissue inhibitor of metalloproteinase. Created with BioRender.com.

#### 3.1.1 Disease initiation mechanism

In the interstitial fibrosis model (Hao et al., 2014), epithelial cell damage is the initiator of fibrosis. However, in diabetes-induced glomerular fibrosis, the consensus is that high glucose initiates fibrosis through mesangial cells, not epithelial cells (Thomas and Ford Versypt, 2022), as collagen is increasingly accumulated when mesangial cells are cultured in high glucose conditions (Ayo et al., 1990, 1991; Haneda et al., 1991; Baccora et al., 2007).

Here, we considered a mechanism in which high glucose initiates fibrosis through mesangial cells via AGE. AGE can cause inflammation within mesangial cells (Yamagishi et al., 2002; Sun et al., 2017) and is implicated in the progression of glomerular damage in diabetes. AGE has been shown to rapidly form when ECM is incubated in high glucose conditions (Singh et al., 2014) and has been observed to readily accumulate within the glomerulus of diabetic mice (Sohn et al., 2013; Do et al., 2018; Iacobini et al., 2018).

Mesangial cells produce monocyte chemoattractant protein (MCP) in the *in vitro* diabetic glomerular fibrosis model (Min et al., 2009) and *in vivo* during diabetes-induced fibrosis (Li et al., 2020). MCP is a chemoattractant commonly associated with the recruitment of macrophages. The inflammation induced by AGE has been shown to enhance MCP expression by mesangial cells (Yamagishi et al., 2002; Sun et al., 2017). The inflammation of mesangial cells and the enhanced production of MCP support the mechanism of MCP-mediated recruitment of macrophages.

#### 3.1.2 Immune cell recruitment

Macrophages are part of the immune response to injury. As such, they can be found across the body during different inflammatory scenarios with cellular damage. Macrophages are known to be recruited to the site of damage via the production of chemoattractants by the damaged cells. Here, we considered macrophage recruitment to the glomerulus through MCP where the inflamed mesangial cells are the source of the MCP. This MCP causes the recruitment and activation of macrophages from precursor monocytes.

The recruitment and accumulation of macrophages within the glomerulus during diabetes are well documented. Many *in vivo* studies in diabetic mice show a significant accumulation of macrophages within the glomerulus a short period after the mice develop the diabetic condition (Ichinose et al., 2006; Saito et al., 2011; Kim et al., 2013; Hong et al., 2014; Terami et al., 2014; Choi et al., 2018; Kim et al., 2018; Hwang et al., 2019).

#### 3.1.3 Mesangial cell activation

In the interstitial fibrosis model, transforming growth factor-β (TGF-β) activates the resident fibroblasts into myofibroblasts. Here, TGF-β is involved in diabetic glomerular fibrosis in a similar capacity. *In vitro* studies where mesangial cells are incubated in high glucose have shown that TGF-β is an essential mediator of fibrosis, as demonstrated by its upregulation in high glucose conditions (Oh et al., 1998; Baccora et al., 2007). Many *in vivo* studies have also shown TGF-β upregulation in diabetic mice (Hong et al., 2001, 2014; Park et al., 2016; Choi et al., 2018; Kim et al., 2018). TGF-β inhibition leads to a decrease in collagen accumulation (Chen et al., 2003). The role of TGF-β as a critical regulator of fibrosis is further supported by the many computational models of fibrosis in other organs and disease cases that have used TGF-β in this role (Sáez et al., 2013; Hao et al., 2014; Rikard et al., 2019; Islam et al., 2023). In other fibrosis scenarios, TGF-β has been implicated in signaling for certain cell types, such as epithelial cells, to transition into a more fibrotic phenotype (Sutariya et al., 2016). The observed TGF-β-mediated increase in collagen in diabetes likely occurs through TGF-β’s activation of the mesangial cells (Liu, 2006; Simonson,2007; Li et al., 2020). Consequently, we included TGF-β as the extracellular signaling molecule involved in fibrosis by activating mesangial cells.

In the interstitial fibrosis model, the source of TGF-β is macrophages. Macrophages produce matrix-metalloproteinase (MMP)9, which is capable of cleaving collagen IV within the ECM. The ECM is embedded with TGF-β that is released when the ECM becomes degraded. Instead of incorporating the entire mechanism for TGF-β release, we kept the simplification that macrophages produce TGF-β, which has the same net effect of macrophages producing MMP that then causes the release of TGF-β.

#### 3.1.4 Collagen production and degradation

Excess production of collagen is seen in *in vitro* studies of mesangial cells incubated in high glucose, a culture model representative of the glomerulus in diabetes (Ayo et al., 1990, 1991; Haneda et al., 1991; Baccora et al., 2007). These studies showed that collagen protein is significantly upregulated in the diabetic milieu. Similarly, *in vivo* studies in spontaneously diabetic mice also showed significant amounts of collagen deposited in the mesangium of the glomerulus after a few weeks of hyperglycemia (Ichinose et al., 2006; Kosugi et al., 2010; Kim et al., 2013; Chen et al., 2014; Hong et al., 2014; Park et al., 2016; Choi et al., 2018; Kim et al., 2018; Hwang et al., 2019). However, the source of the excess collagen protein is not well defined. These *in vitro* and *in vivo* studies do not delineate whether quiescent mesangial cells produce excess collagen or if a different phenotype of mesangial cells produces excess collagen. In other fibrosis mechanisms, fibroblasts (resident cells) get activated to become myofibroblasts, an increasingly fibrotic phenotype that leads to the excess production of collagen (Hao et al., 2014). *In vitro* mesangial cell studies do not explicitly state that the activation of mesangial cells leads to the phenotype that is increasingly fibrotic. However, other studies have made these parallels of mesangial cells becoming activated or differentiating and behaving similarly to myofibroblasts (Simonson, 2007; Li et al., 2020). Mesangial cells have been shown to exhibit similar behaviors to myofibroblasts, such as increased expression of α-smooth muscle actin (α-sma), higher contractility, and increased production of interstitial collagen (Johnson et al., 1992; Liu, 2006; Thomas and Ford Versypt, 2022). Thus, we incorporated the activation of mesangial cells into our model and specified that these cells are the source of the excess collagen.

The mechanism of collagen production and degradation considered here was built upon well-established protein-enzyme interactions. Collagen is a protein degraded by enzyme MMPs, which are inhibited by tissue inhibitor of metalloproteinases (TIMP). These proteins, enzymes, and inhibitors vary in the *in vitro* diabetic culture model (McLennan et al., 1994; Wahab and Mason, 1996; McLennan et al., 2000), indicating that their involvement is pertinent to the progression of glomerular fibrosis.

An *in vitro* co-culture study of mesangial and macrophage cells showed that macrophages regulate MMP expression but not expression of TIMP (Min et al., 2009). In contrast in the same paper, mesangial cells were shown to regulate both MMP and TIMP expression (Min et al., 2009). This differs from the interstitial fibrosis case, where macrophages regulate both MMP and TIMP, and the resident fibroblast cell in that context has no role in MMP and TIMP regulation. Currently, in our model, the sources of MMP and TIMP are the same as in the interstitial fibrosis case. In future iterations of the model, variations in the sources of MMP and TIMP could be incorporated.

### 3.2 Model equations

Here, we describe the equations used to model glomerular fibrosis in diabetes, the specific biological interpretation of each of the terms in the equations, and how the equations for the model were derived.

A mathematical model for interstitial fibrosis in lupus nephritis was built by Hao et al. (2014). Our approach was to adapt their model to glomerular fibrosis in diabetes using the mechanistic steps identified in Section 3.1. In this adapted mathematical model (Figure 1), ten species are the critical cells and biomolecules involved in the process of glomerular fibrosis in diabetes. The ten species consist of three cell types: mesangial cells, activated mesangial cells, and macrophages. The remaining seven species are biomolecules: glucose, AGE, MCP, TGF-β, collagen, MMP, and TIMP.

The dynamics for the species depend on biological processes that lead to changes in their amounts within the glomerulus. The main changes in biomolecule concentrations are due to production by particular cells within the glomerulus and their natural and enzymatic degradation. The main changes in cellular populations are due to cell proliferation, infiltration of immune cells, differentiation of resident cells, and natural death.

These biological processes were modeled using mass action, Michaelis-Menten, or Hill function type kinetics. For biological processes where a biomolecular activates cells to a phenotypic change or a biomolecule stimulates cells to produce another biomolecule, Michaelis-Menten or Hill function type kinetics were used because the stimulation of the cell through a biomolecule interacting with the receptors on the cell is a receptor-limited process. The receptors become saturated. Thus, even at high ligand concentrations, the rate of cellular stimulation by ligands reaches a maximum. The use of Michaelis-Menten or Hill function type kinetics for modeling the stimulation and activation of cells is a common approach (Hao et al., 2014, 2015; Ruggiero et al., 2017; Islam et al., 2021; Cook et al., 2022; Patidar and Ford Versypt, 2023; Islam et al., 2023; Cook et al., 2024). Equation (1) defines the generalized Hill function for activation by any species X as

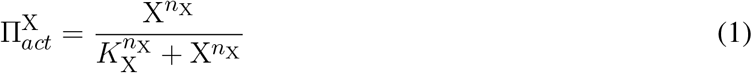

where *K*_X_ and *n*_X_ represent the saturation constant and Hill function parameter, respectively. Michaelis-Mention kinetics result when *n*_X_ = 1.

Using these modeling approaches and adapting equations from Hao et al. (2014), the following set of equations were defined to model each of the steps within the progression of glomerular fibrosis in diabetes (Figure 1).

#### 3.2.1 Disease initiation

The progression of glomerular fibrosis in diabetes is initiated by glucose stimulating the formation of AGE. Equation (2) defines the resulting AGE dynamics as

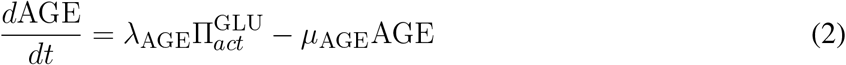

where the first term represents the rate of production of AGE dictated by the AGE formation rate constant λ_AGE_ and the 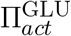 Hill function for activation by glucose with parameters *K*_GLU_ and *n*_GLU_. The second term describes the first-order AGE removal rate, which depends on the degradation rate constant *μ*_AGE_.

Equation (3) describes the dynamics of MCP, the chemoattractant protein that is involved in the recruitment of macrophages, as

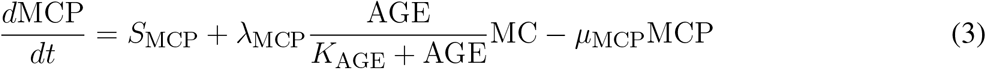

where *S*_MCP_ is the baseline production of MCP. The second term is the AGE-induced MCP production by mesangial cells (MC) modeled using Michaelis-Menten kinetics (Equation (1)) where λ_MCP_ and *K*_AGE_ represent the production rate and saturation constant, respectively. The last term represents the MCP removal rate from the glomerulus with first-order rate constant *μ*_MCP_.

The population of mesangial cells (MC) is assumed to be constant at the value listed in Table 1. Activated mesangial cells (AMC) are considered as a separate population.

**Table 1.** Parameters for glomerular fibrosis model. Abbreviations: AGE, advanced glycation end products; GLU, glucose; MCP, monocyte chemoattractant protein; MC, mesangial cells; MAC, macrophages; TGF, transforming growth factor–β; AMC, activated mesangial cells; MMP, matrix metalloproteinase; TIMP, tissue inhibitor of metalloproteinase; COL, collagen.

| Parameter | Symbol | Value | Units | Source |
| --- | --- | --- | --- | --- |
| Production rate of AGE | $\lambda_{AGE}$ | $1.54 \times 10^{-5}$ | $(\text{g/ml})^{-5} \text{ day}^{-1}$ | Estimated |
| GLU saturation constant for AGE production | $K_{GLU}$ | $3.37 \times 10^{-2}$ | g/ml | Estimated |
| Hill function power term for AGE production by GLU | $n_{GLU}$ | 2.7 | Unitless | Estimated |
| AGE degradation rate | $\mu_{AGE}$ | $8.7 \times 10^{-3}$ | $\text{day}^{-1}$ | Rabbani and Thornalley (2018) and estimated |
| Baseline MCP production rate | $S_{MCP}$ | $2.77 \times 10^{-10}$ | $(\text{g/ml}) \text{ day}^{-1}$ | Estimated |
| Production rate of MCP by MC | $\lambda_{MCP}$ | $4.09 \times 10^{-10}$ | $\text{day}^{-1}$ | Estimated |
| AGE saturation constant for MCP production | $K_{AGE}$ | $1.14 \times 10^{-6}$ | g/ml | Estimated |
| MCP degradation rate | $\mu_{MCP}$ | 1.73 | $\text{day}^{-1}$ | Hao et al. (2014) |
| Baseline MCP concentration *in vitro* | $MCP_{\text{vitro}}$ | $1.60 \times 10^{-10}$ | g/ml | Yamagishi et al. (2002) |
| Recruitment rate for MAC | $\lambda_{MAC}$ | $6.2 \times 10^{-2}$ | $\text{day}^{-1}$ | Estimated |
| MCP saturation constant for MAC recruitment | $K_{MCP}$ | $5.00 \times 10^{-9}$ | g/ml | Hao et al. (2014) |
| Hill function power term for MAC recruitment by MCP | $n_{MCP}$ | 5.38 | Unitless | Estimated |
| Constant macrophage density in the blood | $MAC_0$ | $5.00 \times 10^{-5}$ | g/ml | Hao et al. (2014) |
| MAC death/removal rate | $\mu_{MAC}$ | 0.15 | $\text{day}^{-1}$ | Estimated |

Table 1 continued from previous page
| Parameter | Symbol | Value | Units | Source |
| --- | --- | --- | --- | --- |
| Baseline TGF production rate | $S_{\text{TGF}}$ | $2.85 \times 10^{-7}$ | (g/ml) day<sup>-1</sup> | Estimated |
| Production rate of TGF by MAC | $\lambda_{\text{TGF}}$ | $1.34 \times 10^5$ | day<sup>-1</sup> | Estimated |
| Degradation rate of TGF | $\mu_{\text{TGF}}$ | $3.33 \times 10^2$ | g/ml | Hao et al. (2014) |
| Baseline AMC activation rate | $S_{\text{AMC}}$ | $2.6 \times 10^{-4}$ | (g/ml) day<sup>-1</sup> | Estimated |
| TGF-β mediated AMC activation rate | $\lambda_{\text{AMC}}$ | $4 \times 10^{-3}$ | day<sup>-1</sup> | Estimated |
| TGF-β saturation constant for activation of MC | $K_{\text{TGF}}$ | $2.5 \times 10^{-9}$ | g/ml | Estimated |
| Hill function power term for MC activation by TGF | $n_{\text{TGF}}$ | 4.14 | unitless | Estimated |
| AMC death rate *in vitro* | $\mu_{\text{AMC}_{\text{vitro}}}$ | 0.0166 | day<sup>-1</sup> | Estimated |
| AMC death rate | $\mu_{\text{AMC}}$ | $5 \times 10^{-1}$ | day<sup>-1</sup> | Estimated |
| Production rate of MMP by MAC | $\lambda_{\text{MMP}}$ | $2.06 \times 10^7$ | day<sup>-1</sup> | Estimated |
| Binding rate of MMP to TIMP in molar units | $\gamma$ | $1.6 \times 10^5$ | M<sup>-1</sup>s<sup>-1</sup> | Vempati et al. (2007) |
| Binding rate of MMP to TIMP ($\gamma/\text{MW}_{\text{MMP}}$) | $\gamma_{\text{MMP}}$ | $1.46 \times 10^8$ | (g/ml)<sup>-1</sup> day<sup>-1</sup> | Vempati et al. (2007) and estimated |
| Degradation rate of MMP | $\mu_{\text{MMP}}$ | 4.32 | day<sup>-1</sup> | Hao et al. (2014) |
| Production rate of TIMP by MAC | $\lambda_{\text{TIMP}}$ | $6.00 \times 10^{-9}$ | day<sup>-1</sup> | Estimated |
| Binding rate of TIMP to MMP ($\gamma/\text{MW}_{\text{TIMP}}$) | $\gamma_{\text{TIMP}}$ | $0.69 \times 10^9$ | (g/ml)<sup>-1</sup> d<sup>-1</sup> | Vempati et al. (2007) and estimated |
| Degradation rate of TIMP | $\mu_{\text{TIMP}}$ | 21.60 | day<sup>-1</sup> | Hao et al. (2014) |
| Production rate of COL by MC | $\lambda_{\text{COL}}$ | $3.00 \times 10^{-3}$ | day<sup>-1</sup> | Hao et al. (2014) |
| Production rate of COL by AMC | $\lambda_{\text{COLA}}$ | $1.83 \times 10^3$ | day<sup>-1</sup> | Estimated |
| Degradation rate of COL | $\mu_{\text{COL}}$ | $3.70 \times 10^{-1}$ | day<sup>-1</sup> | Hao et al. (2014) |
| Degradation rate of COL by MMP | $\gamma_{\text{COL}}$ | $1.84 \times 10^4$ | (g/ml)<sup>-1</sup> d<sup>-1</sup> | Estimated |
| Blood GLU concentration baseline | $G_1$ | $1 \times 10^{-3}$ | (g/ml) | Averaged value from Figure 3 |
| Blood GLU concentration diabetic | $G_2$ | $5 \times 10^{-3}$ | (g/ml) | Averaged value from Figure 3 |
| Start time for blood GLU linear increase | $t_1$ | 6 | weeks | Extracted value from Figure 3 |

Table 1 continued from previous page
| Parameter | Symbol | Value | Units | Source |
| --- | --- | --- | --- | --- |
| End time for blood GLU linear increase | $t_2$ | 16 | weeks | Extracted value from Figure 3 |
| Inhibition term for treatment scenarios | $K_I$ | $1 \times 10^5$ | - | Set value |

#### 3.2.2 Immune cell recruitment

Equation (4) defines the population dynamics of the macrophages (MAC) recruited by MCP as

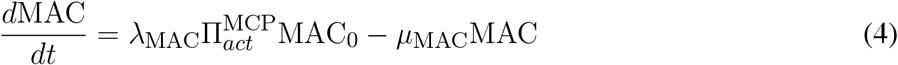

where the first term represents the MCP-dependent recruitment of macrophages from the blood dictated by the maximum recruitment rate λ_MAC_, the 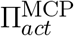 Hill function for activation by MCP (Equation (1)), and the constant macrophage density in the blood MAC_0_. The second term in Equation (4) represents the removal of macrophages from the glomerulus due to death and phenotypical switch with a combined rate constant *μ*_MAC_.

#### 3.2.3 Mesangial cell activation

Equation (5) defines the dynamics for TGF-β (TGF), the growth factor involved in the activation of mesangial cells, as

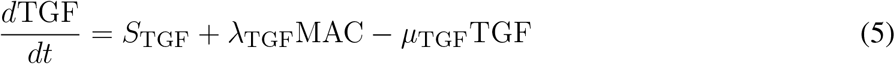

where the first two terms represent the baseline production of TGF-β and the macrophage-mediated production of TGF-β having the production rate constants *S*_TGF_ and λ_TGF_, respectively. The natural degradation of TGF-β is represented by the last term with a degradation rate constant *μ*_TGF_.

Equation (6) defines the population dynamics for activated mesangial cells (AMC) as

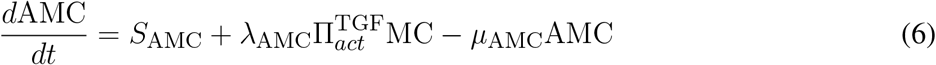

where λ_AMC_ is the activation rate constant and 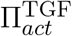 is the Hill function for the TGF-β mediated activation of mesangial cells (Equation (1)). The last term represents the death or apoptosis of AMC with a rate constant *μ*_AMC_.

#### 3.2.4 Collagen production and degradation

Collagen degradation depends on the enzymes MMP and TIMP. As in Hao et al. (2014), the population dynamics of MMP and TIMP are defined by Equations (7) and (8), respectively.

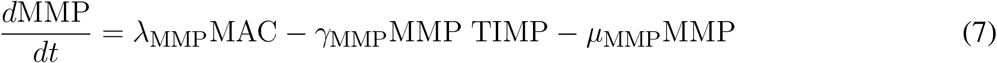

The first, second, and third terms of Equation (7) represent macrophage-mediated production, MMP inhibition by TIMP, and degradation, respectively. The terms have rate constants λ_MMP_, *γ*_MMP_, and *μ*_MMP_, respectively.

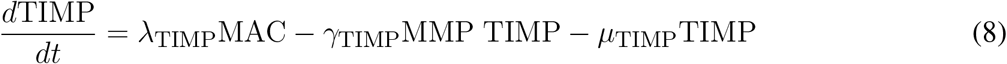

The first, second, and third terms of Equation (8) represent macrophage-mediated production, TIMP binding with MMP, and the degradation of TIMP, respectively. The terms have rate constants λ_TIMP_, *γ*_TIMP_, and *μ*_TIMP_, respectively. Note that *γ*_MMP_ and *γ*_TIMP_ only differ by unit conversions from a molar basis to the mass per volume basis for MMP and TIMP, respectively (Table 1).

Equation (9) defines the dynamics of collagen (COL) as

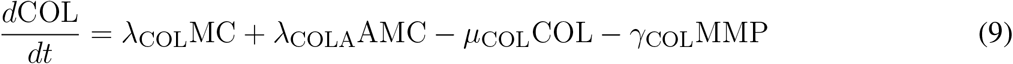

where the first two terms represent collagen production by MC and AMC, respectively. The last two terms represent the natural degradation of collagen and the MMP-mediated degradation of collagen. Each of the terms are dictated by corresponding rate constants λ_COL_, λ_COLA_, *μ*_COL_, and *γ*_COL_.

There are several differences between the Hao et al. (2014) model for interstitial fibrosis and our model beyond the application to glomerular fibrosis. Foremost, our model does not have a spatial component for the transport in the small mesangial region for glomerular fibrosis. Equations (2) and (3) were newly included here because of mechanistic differences between glomerular and interstitial fibrosis. Equation (4) has a similar mechanism as the interstitial model, but due to the removal of the spatial component, the recruitment and activation of macrophages term was modeled using a simpler Hill function than the boundary condition-dependent equation implemented in Hao et al. (2014). Equation (5) has a baseline TGF-β production term added to better fit the data. Equation (6) has only a TGF-β-mediated activation term instead of activation via TGF-β and platelet-derived growth factor. The reaction portions of Equations (7) and (8) were left unchanged due to mechanistic similarities. Equation (9) had a TGF-β-mediated collagen production term removed from the Hao et al. (2014) model since TGF-β was already playing a significant role in the accumulation of collagen through the activation of mesangial cells, which is the source of the excess collagen.

### 3.3 Parameter estimation

The model parameters were calibrated based on a combination of *in vivo* and *in vitro* data and parameter values obtained from the literature (Section 1 and Figure 2). Parameters were estimated using a subsystem approach where only three to four parameters were determined simultaneously. The estimation was divided such that the parameters involved in the dynamics of one group of species were estimated before moving on to the next. The process was continued until all parameters were calibrated. As such, we discuss the estimation of parameters involving each group of species separately below. Unless stated otherwise, nonlinear least squares regression in MATLAB was used to minimize the sum of squared differences between model predictions and experimental data. The resulting parameter values are listed in Table 1.

**Figure 2.**
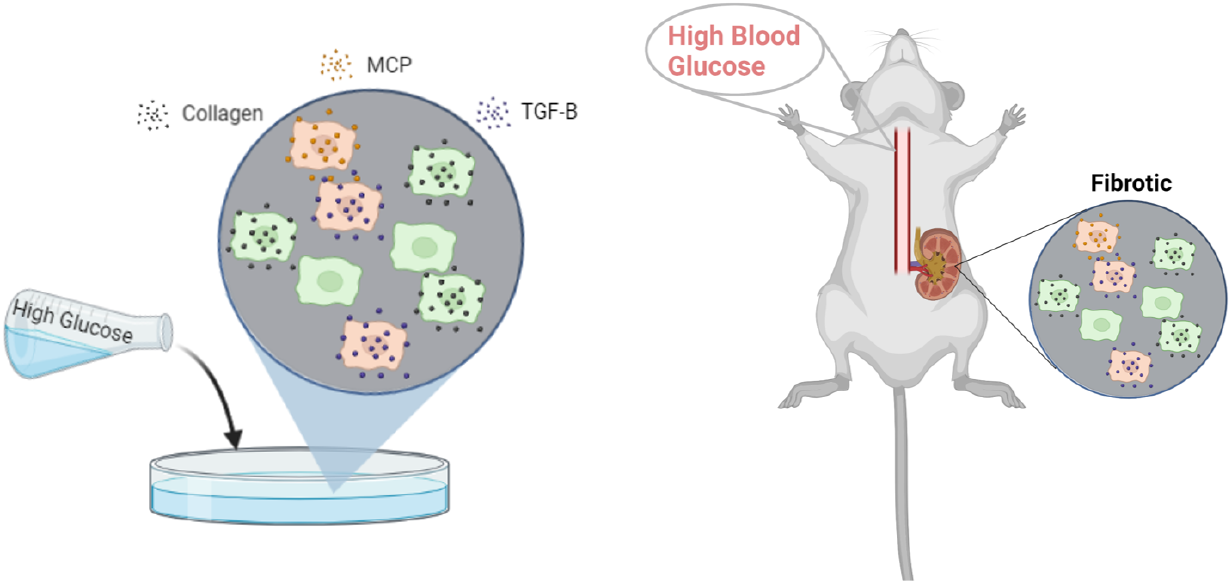
Published *in vitro* mesangial cell culture and *in vivo* db/db mice glomerular data were used for parameter estimation. Abbreviations: MCP, monocyte chemoattractant protein; TGF-β, transforming growth factor–β. Created with BioRender.com.

#### 3.3.1 Glucose and AGE

The glomerular fibrosis model defined in Section 3.2 uses an input of blood glucose concentration as the stimulus for fibrosis progression (Figure 3A). For this input, we used the blood glucose concentration of db/db mice over 24 weeks (Figures S1 and 3B) (Cohen et al., 1995, 2001, 2002; Ziyadeh et al., 2000; Koya et al., 2000; Kolavennu et al., 2008). We approximated the data as a ramp input to simplify the model. The input GLU (Equation (10)) is fed into the model via the AGE dynamics (Equation (2)).

**Figure 3.**
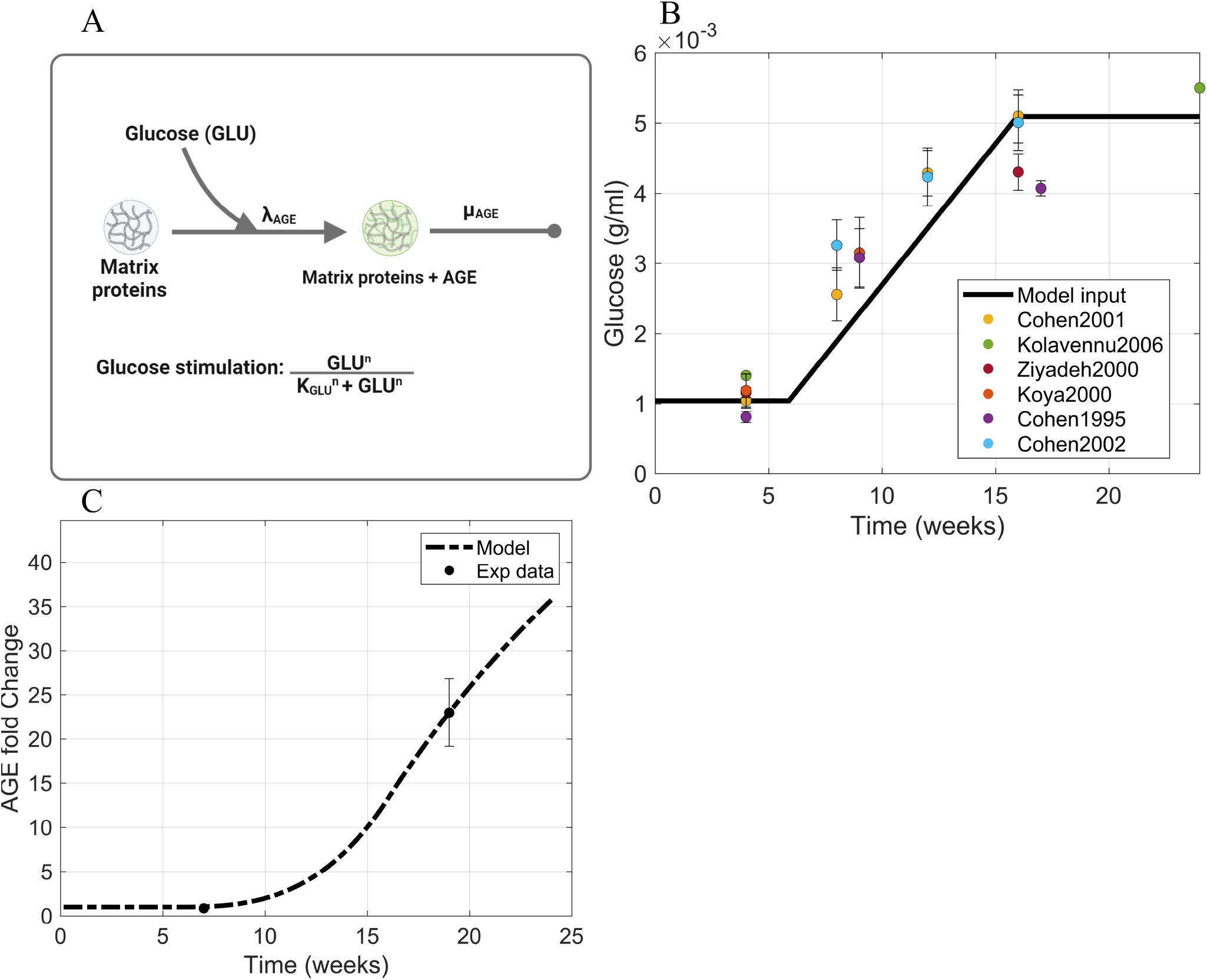
Information supporting AGE parameter estimation. (A) The rate constants and species contributing to AGE dynamics (Equation (2)). (B) Blood glucose concentration dynamics within db/db mice approximated as a piecewise function of ramp input between two constant intervals. Sources: Cohen et al. (1995, 2001, 2002); Ziyadeh et al. (2000); Koya et al. (2000); Kolavennu et al. (2008). (C) Fitting of AGE time-series data to estimate parameters involved in AGE dynamics. Source: Sohn et al. (2013). Raw data points are shown in Figure S1. Abbreviations: AGE, advanced glycation end products; GLU, glucose.

Equation (10) defines the dynamics of the glucose (GLU) concentration as a piecewise function

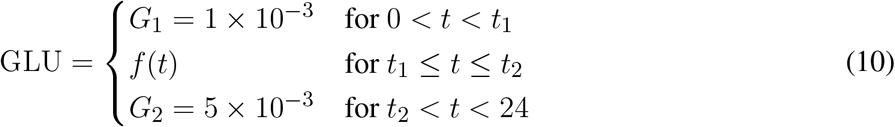

where *t* is in time in weeks, and *f* (*t*) is a linear function defined as

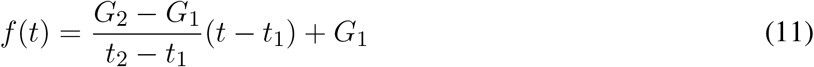

that approximates the increase in glucose concentration in the interval of [*t*_1_, *t*_2_] = 6–16 weeks. In Equation (11), (*G*_1_, *t*_1_) and (*G*_2_, *t*_2_) are the starting and ending coordinates for the interval of linear increase of glucose concentration between two intervals constant at *G*_1_ and *G*_2_. The values are listed in Table 1.

AGE is formed through the non-enzymatic reaction of glucose with proteins to form a covalently bonded, stable molecule. Different types of AGE can be formed due to high glucose concentration within the body, such as glycated albumin, hydroimidazolones, and glucosepane (Rabbani and Thornalley, 2018). Out of the many types of AGE, Nϵ-(carboxymethyl) lysine (CML) is expressed in the highest quantities in mice (Kim et al., 2018; Menini et al., 2018) relative to other AGE (Doan et al., 2015; Kim et al., 2018). We assumed that CML has the highest impact on the progression of fibrosis in diabetes; thus, we based the AGE dynamics on CML.

We calculated the AGE degradation rate *μ*_AGE_ using the half-life of collagen *t*_1*/*2,COL_ ≈ 80 days (Brings et al., 2017) via *μ*_AGE_ = ln(2)*/t*_1*/*2,COL_. We used the half-life of collagen instead of CML because CML can be enzymatically removed when collagen turnover occurs, which happens at a much faster rate than the degradation of CML (Rabbani and Thornalley, 2018).

The AGE initial value AGE_*ss*_ was assumed to be the CML baseline serum concentration obtained from mice and rats (Yuan et al., 2017; Menini et al., 2018). The AGE rate of formation λ_AGE_ and the Hill function power term *n*_GLU_ were estimated from AGE concentration fold change experimental data obtained from the glomeruli of db/db mice (Figure 3C) (Sohn et al., 2013). In this way, the parameters involved in AGE dynamics were estimated, and the resulting values are in Table 1.

#### 3.3.2 MCP

To calibrate the Michaelis-Menten parameters λ_MCP_ and *K*_AGE_ and the baseline MCP production rate MCP_0_ in Equation (3) (Figure 4A), data for *in vitro* expression of MCP by AGE-stimulated mesangial cells were used (Figure 4B) (Lu et al., 2004).

**Figure 4.**
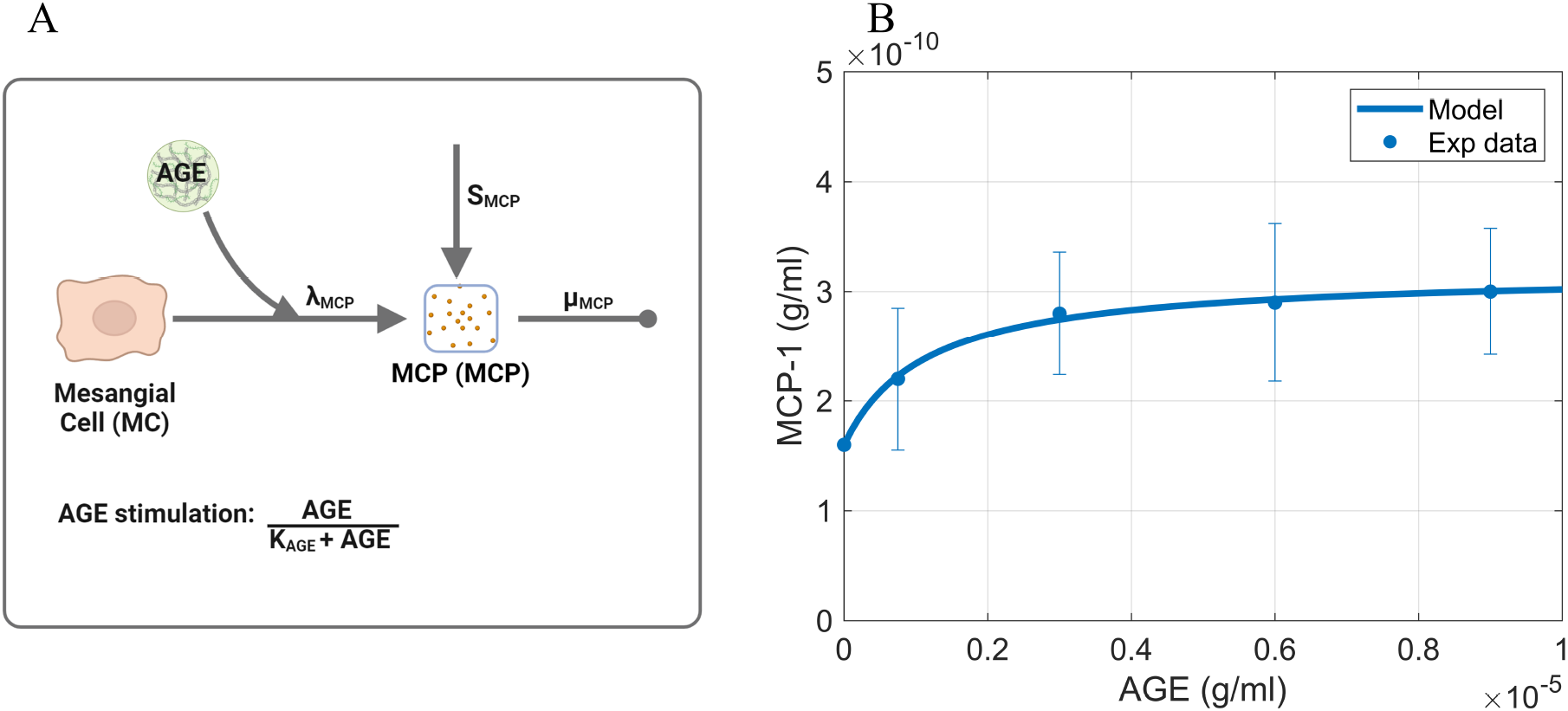
Information supporting MCP parameter estimation. (A) The rate constants and species contributing to the dynamics of MCP (Equation (3)). (B) Fitting of dose-response curve to estimate Michaelis-Menten parameters. Source: Lu et al. (2004). Raw data points are shown in Figure S2. Panel A created with BioRender.com. Abbreviations: AGE, advanced glycation end products; MCP, monocyte chemoattractant protein; MC, mesangial cells.

The baseline MCP production rate *S*_MCP_ was calculated by setting Equation (3) to steady state and the AGE concentration stimulus to zero to obtain

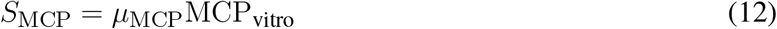

where *μ*_MCP_ is the degradation rate of MCP from the literature (Hao et al., 2014) and MCP_vitro_ is the baseline concentration of MCP in the absence of AGE stimulation, which was obtained from the processed *in vitro* MCP expression data (Figures S2 and 4B). For estimating the Michaelis-Menten parameters λ_MCP_ and *K*_AGE_, we used the values of *S*_MCP_ and *μ*_MCP_ and the *in vitro* MCP expression data (Figure 4B) (Lu et al., 2004). The acquired values are shown in Table 1.

#### 3.3.3 Macrophages

The macrophage dynamics (Equation (4)) are dictated by the macrophage recruitment rate constant λ_MAC_, the saturation constant for the MCP mediated recruitment of macrophages *K*_MCP_, the macrophage density in the blood MAC_0_, and the death/removal rate of macrophages from the glomerulus *μ*_MAC_ (Figure 5A).

**Figure 5.**
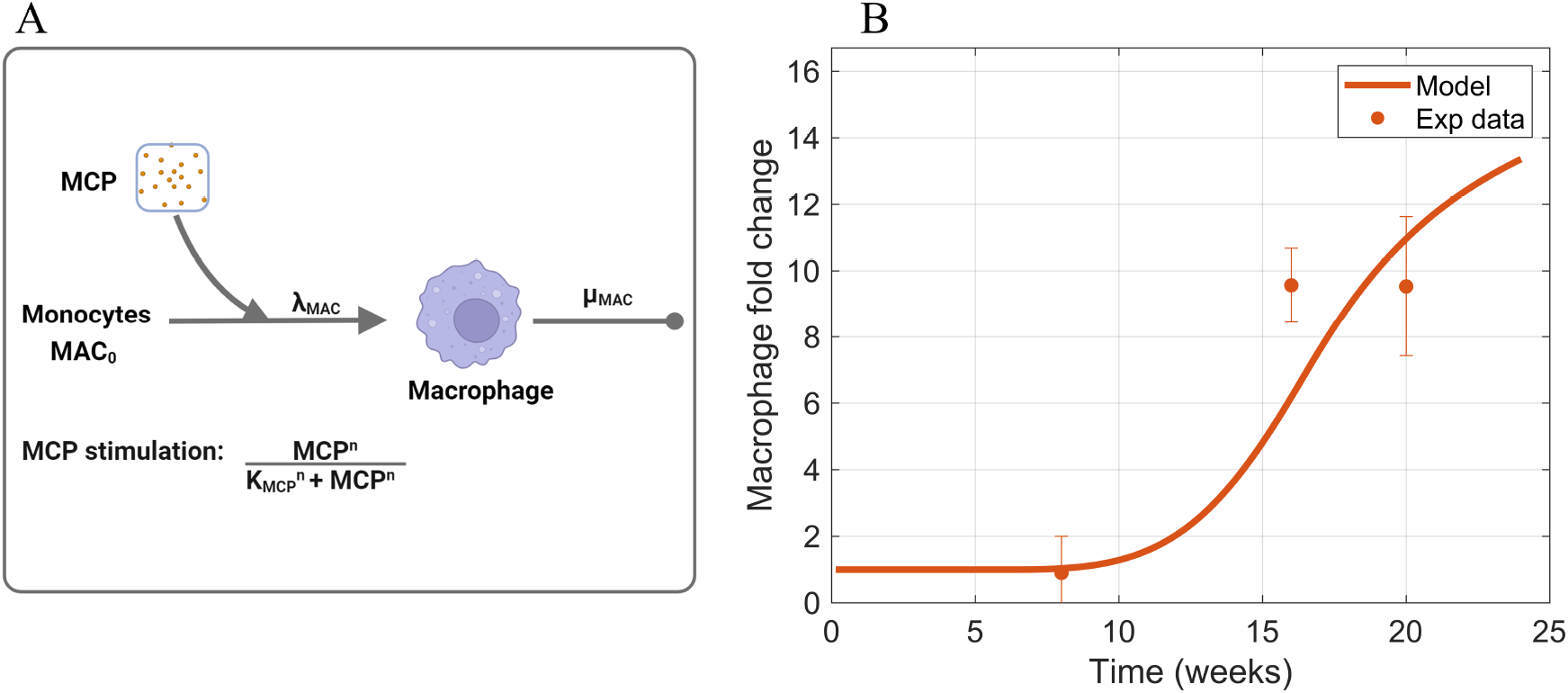
Information supporting MAC parameter estimation. (A) The rate constants and species contributing to the dynamics of macrophages (Equation (4)). (B) Fitting of macrophage time-series data to estimate parameters that dictate macrophage dynamics. Sources: Ichinose et al. (2006); Saito et al. (2011); Kim et al. (2013); Hong et al. (2014); Choi et al. (2018); Kim et al. (2018); Hwang et al. (2019). Non-averaged experimental data points are shown in Figure S3. Panel A created with BioRender.com. Abbreviations: MCP, monocyte chemoattractant protein; MAC, macrophages.

The Hill activation function parameter *K*_MCP_ was obtained from the literature (Hao et al., 2014). The Hill function power term *n*_MCP_, macrophage death rate *μ*_MAC_, and recruitment rate λ_MAC_ were estimated such that the model prediction of macrophage dynamics fit the macrophage time-series data (Figure 5B). The acquired values are shown in Table 1.

#### 3.3.4 TGF-β

TGF-β dynamics (Equation (5)) are dictated by the rate of TGF-β production by macrophages λ_TGF_, the baseline TGF-β production rate *S*_TGF_, and the degradation rate of TGF-β *μ*_TGF_ (Figure 6A). While *μ*_TGF_ was obtained from the literature (Hao et al., 2014), the baseline and macrophage-mediated production rate constants *S*_TGF_ and λ_TGF_, respectively, were estimated using the TGF-β fold change time-series data (Figures S4 and 6B) (Hong et al., 2014; Park et al., 2016; Choi et al., 2018; Kim et al., 2018).

**Figure 6.**
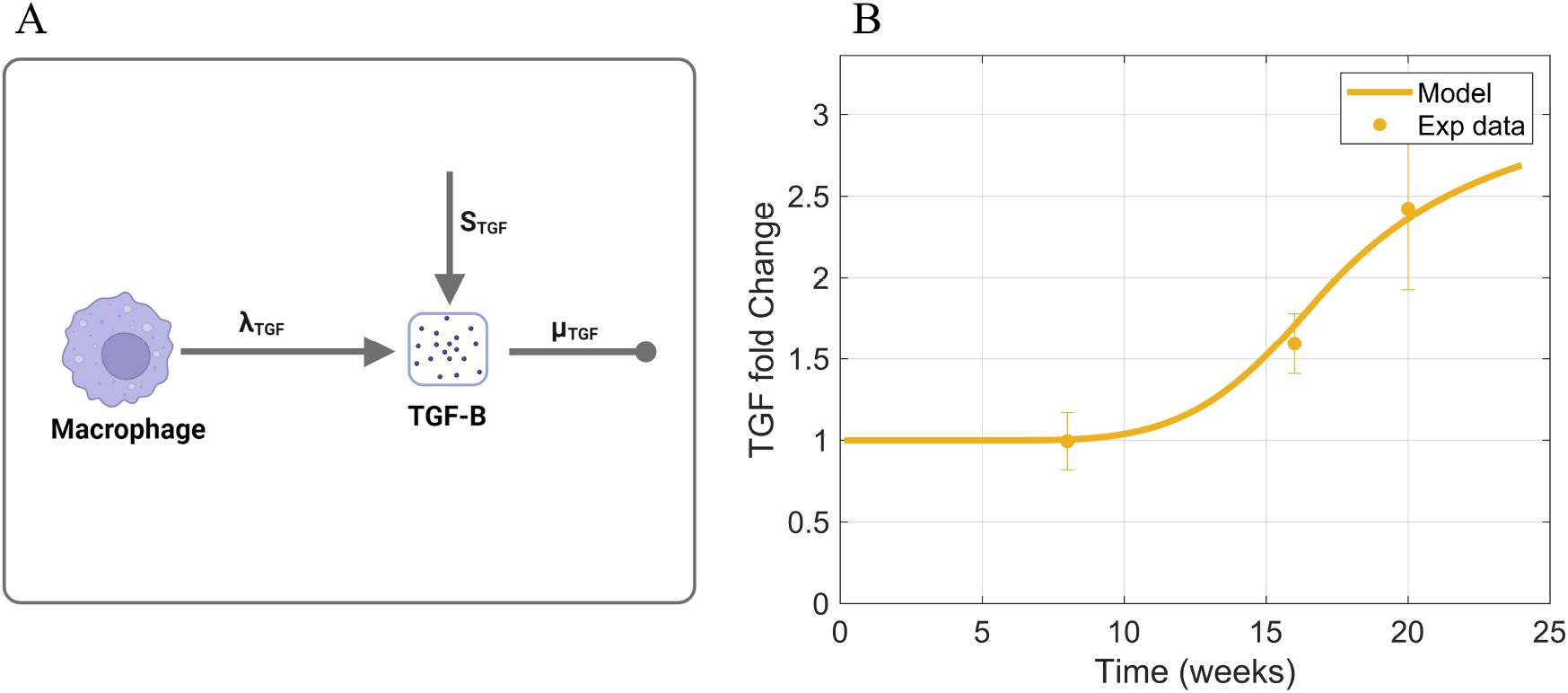
Information supporting TGF-β parameter estimation. (A) The rate constants and species contributing to the dynamics of TGF-β (Equation (5)). (B) Fitting of TGF-β time-series data to estimate parameters that dictate TGF-β dynamics. Sources: Hong et al. (2014); Park et al. (2016); Choi et al. (2018); Kim et al. (2018). Non-averaged experimental data points are shown in Figure S4. Panel A created with BioRender.com. Abbreviations: TGF, transforming growth factor–β.

#### 3.3.5 Activated mesangial cells

Activated mesangial cell dynamics (Equation (6)) depend on a baseline activation of mesangial cells *S*_AMC_, TGF-β mediated activation λ_AMC_, the Hill activation function parameter *K*_TGF_, the Hill function power term *n*_TGF_, and death/removal of activated mesangial cells from the glomerulus *μ*_AMC_ (Figure 7A).

**Figure 7.**
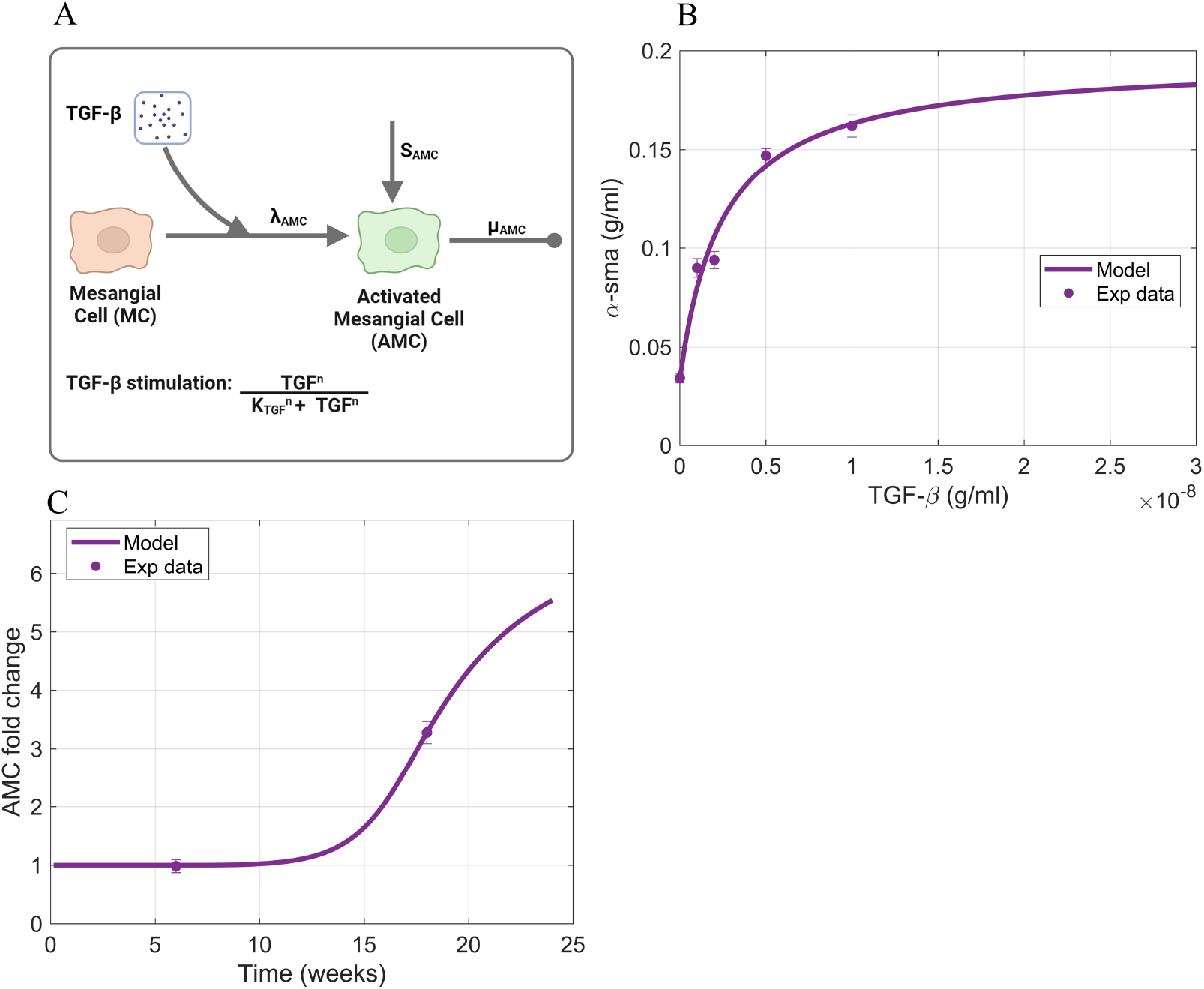
Information supporting AMC parameter estimation. (A) The rate constants and species contributing to the dynamics of activated mesangial cells (Equation (6)). (B) Fitting of TGF-β induced mesangial cell activation as indicated by α-sma expression. Source: Fu et al. (2013). (C) Fitting of activated mesangial cell time-series data to estimate parameters that dictate activated mesangial cell dynamics. Source: Wang et al. (2016). Raw data points are shown in Figure S5. Panel A created with BioRender.com. Abbreviations: MC, mesangial cells; TGF, transforming growth factor–β; AMC, activated mesangial cells.

To estimate the rate parameters involved in each process, we used a combination of *in vitro* and *in vivo* data. The *in vitro* data is the expression of α-sma by TGF-β stimulated mesangial cells, and the *in vivo* data is time-series α-sma fold change data within the glomerulus used as a surrogate for activated mesangial cells (see Section 1.5 for details). The *in vitro* data (Figures S5 and 7B) (Fu et al., 2013) was used to calibrate the mesangial cell activation rate constant λ_AMC_ and the saturation constant for the TGF-β mediated activation of mesangial cells *K*_TGF_. The baseline mesangial cell activation rate constant *S*_AMC_ and the Hill function parameter *n*_TGF_ were estimated from the *in vivo* time-series data from activated mesangial cells (Figures S5 and 7C) (Wang et al., 2016). The death rate of activated mesangial cells 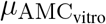 was acquired from literature (Hao et al., 2014). The acquired values are shown in Table 1.

#### 3.3.6 MMP, TIMP, and collagen

MMP and TIMP dynamics (Equations (7) and (8)) are dependent on their production rate constants λ_MMP_ and λ_TIMP_, degradation rate constants *μ*_MMP_ and *μ*_TIMP_, and binding rate constants *γ*_MMP_ and *γ*_TIMP_ (Figure 8A). The MMP production rate constant λ_MMP_ was fitted to steady-state value of MMP obtained from literature, and λ_TIMP_ was constrained to be one-fifth of λ_MMP_, as was done previously in the interstitial fibrosis model (Hao et al., 2014). Degradation rate constants and binding rate constants of MMP and TIMP were gathered from the literature.

**Figure 8.**
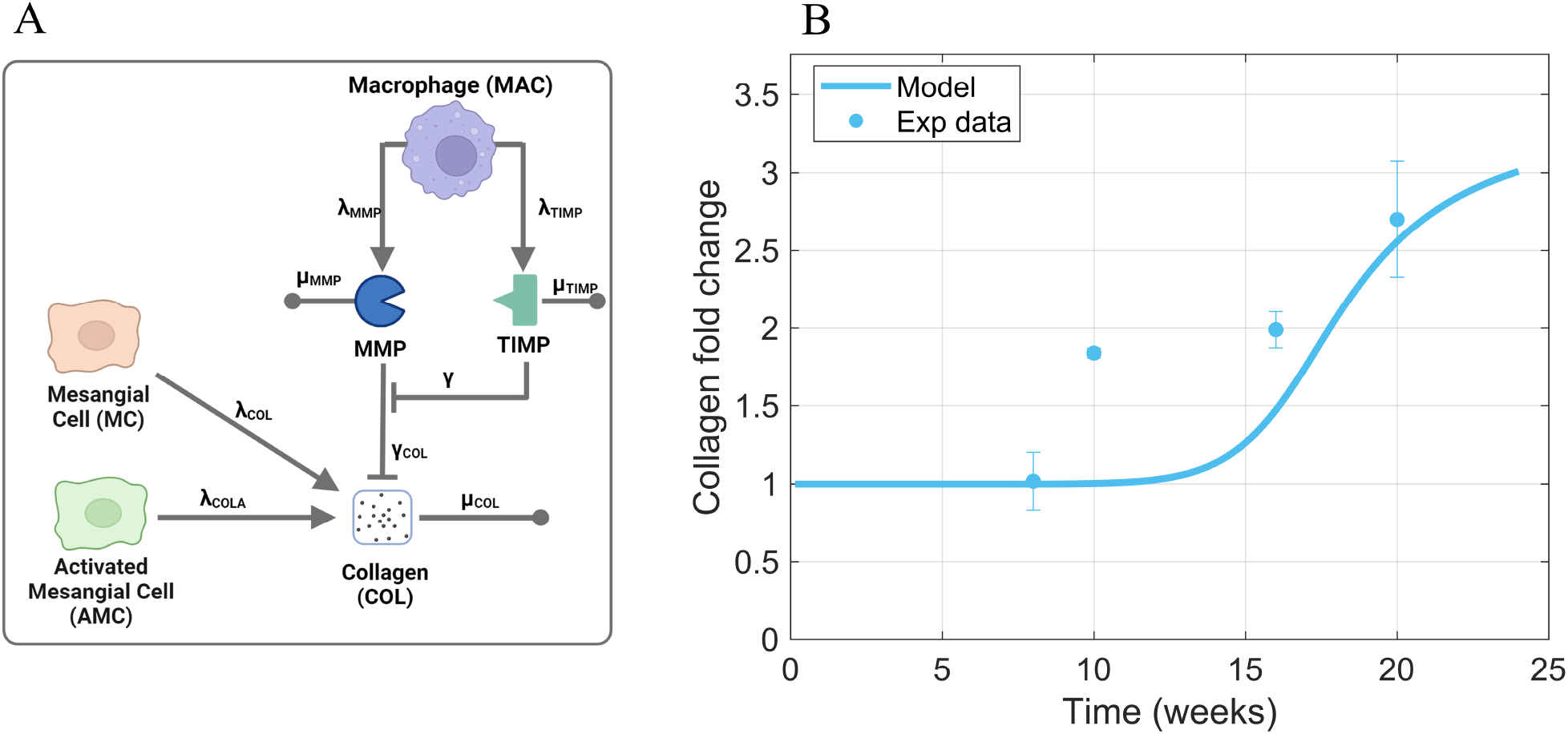
Information supporting MMP, TIMP, and COL parameter estimation. (A) The rate constants and species contributing to MMP, TIMP, and collagen dynamics (Equations (7)–(9)) (B) Fitting of collagen time-series data to estimate parameters that dictate collagen dynamics. Sources: Park et al. (2014); Wang et al. (2016); Nagai et al. (2019); Li et al. (2020); Hsu et al. (2021). Non-averaged data points are shown in Figure S6. Panel A created with BioRender.com. Abbreviations: MC, mesangial cells; MAC, macrophages; TGF, transforming growth factor–β; AMC, activated mesangial cells; MMP, matrix metalloproteinase; TIMP, tissue inhibitor of metalloproteinase; COL, collagen.

Collagen dynamics (Equation (9)) depend on production rate constants λ_COLA_ and λ_COL_ and degradation rate constants *γ*_COL_ and *μ*_COL_ (Figure 8A). To constrain the collagen dynamics to experimental data for collagen protein expression fold change, we fitted the parameter values for the rate of activated mesangial cell-mediated collagen production λ_COLA_ and the MMP-mediated collagen degradation rate *γ*_COL_(Figures S6 and 8B) (Park et al., 2014; Wang et al., 2016; Nagai et al., 2019; Li et al., 2020; Hsu et al., 2021). The collagen production rate constant λ_COL_ and collagen degradation rate constant *γ*_COL_ were obtained from the literature (Hao et al., 2014). The acquired values are shown in Table 1.

### 3.4 Determination of initial values

The initial value for AGE was obtained from the experimental data found in the literature Yuan et al. (2017); Menini et al. (2018). The initial value for AGE within the glomerulus was assumed to be the same as the average serum concentration of CML within healthy mice and rats. The initial values for the other species within the model were estimated by running the model with healthy glucose levels until a steady state was reached for each of the species. Thus, the initial values are denoted with the subscript *ss* for steady state. The gathered and estimated initial values are in Table 2.

**Table 2.**
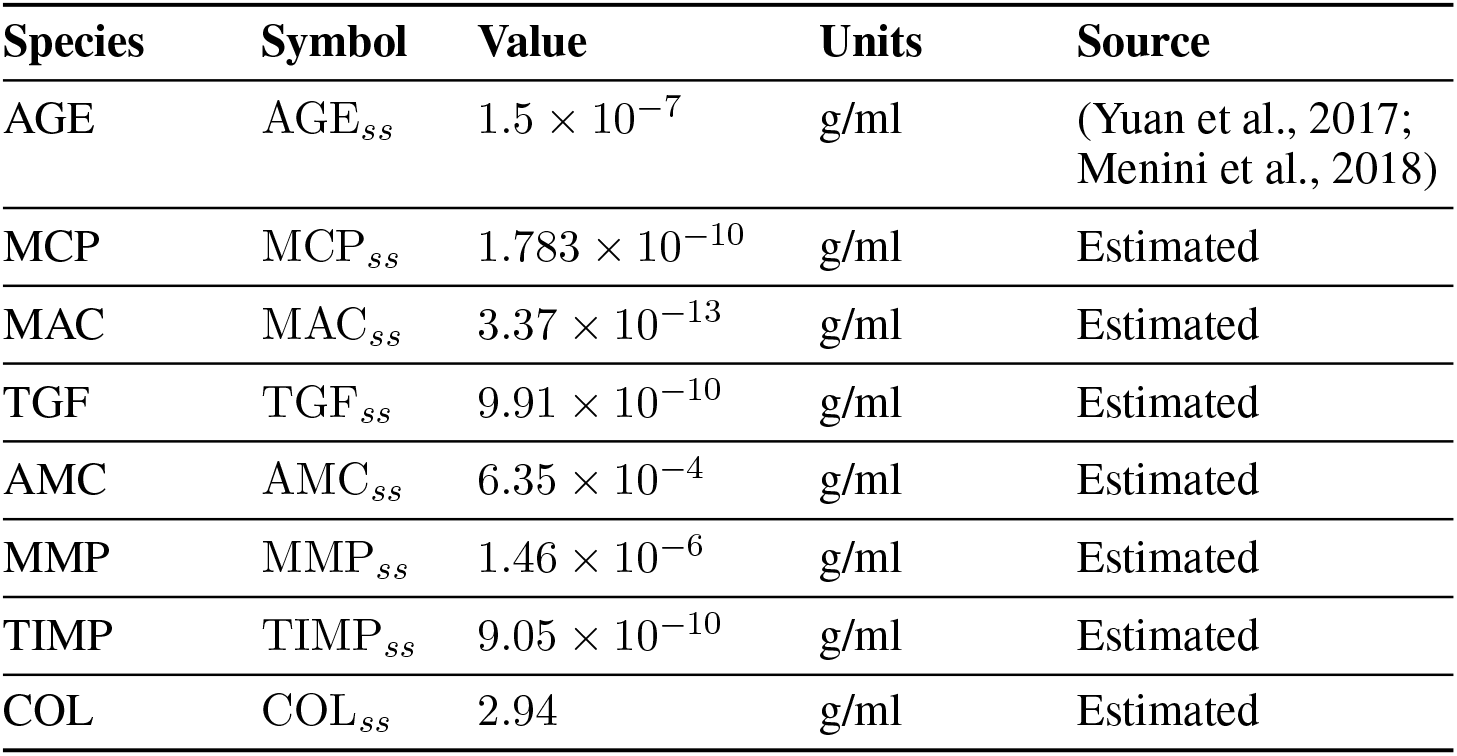
Initial values for model species. Abbreviations: AGE, advanced glycation end products; MCP, monocyte chemoattractant protein; MAC, macrophages; TGF, transforming growth factor–β; AMC, activated mesangial cells; MMP, matrix metalloproteinase; TIMP, tissue inhibitor of metalloproteinase; COL, collagen.

| Species | Symbol | Value | Units | Source |
| --- | --- | --- | --- | --- |
| AGE | $AGE_{ss}$ | $1.5 \times 10^{-7}$ | g/ml | (Yuan et al., 2017; Menini et al., 2018) |
| MCP | $MCP_{ss}$ | $1.783 \times 10^{-10}$ | g/ml | Estimated |
| MAC | $MAC_{ss}$ | $3.37 \times 10^{-13}$ | g/ml | Estimated |
| TGF | $TGF_{ss}$ | $9.91 \times 10^{-10}$ | g/ml | Estimated |
| AMC | $AMC_{ss}$ | $6.35 \times 10^{-4}$ | g/ml | Estimated |
| MMP | $MMP_{ss}$ | $1.46 \times 10^{-6}$ | g/ml | Estimated |
| TIMP | $TIMP_{ss}$ | $9.05 \times 10^{-10}$ | g/ml | Estimated |
| COL | $COL_{ss}$ | 2.94 | g/ml | Estimated |

## 4 RESULTS

We simulated different scenarios to answer key questions that were raised in the introduction. We first simulated the base case scenario for high glucose-induced glomerular fibrosis to capture diabetes-induced glomerular fibrosis. Next, we simulated glucose control to identify why glucose control takes years to recover from kidney damage. Then, we simulated different treatment approaches to identify ways to accelerate the recovery from kidney damage. Additionally, we performed a local sensitivity analysis to identify the most influential parameters in the model (Section 2 and Figure S7).

### 4.1 Glomerular fibrosis base case scenario

The base case scenario for glomerular fibrosis is when the model uses a prescribed blood glucose concentration (Equation (10)), specific initial concentrations of cellular and biomolecular species (Table 2), and parameter values based on experimental data for species dynamics within db/db mice or mesangial cell cultures as described in Section 3.3 and Table 1. The base case scenario shows the dynamics of the critical cells and biomolecules involved in the glomerular fibrosis of diabetic mice over 24 weeks (Figure 9). In the base case scenario, the blood glucose concentration increased at six weeks and plateaued at 16 weeks (Figure 9A), causing an increased accumulation of AGE (Figure 9B). This started the cascade of immune cell recruitment followed by mesangial cell activation and eventual collagen accumulation. All species reached a steady state within 24 weeks except for AGE. The AGE concentration did not reach a steady state within the given time frame because its degradation rate was small relative to its formation rate.

**Figure 9.**
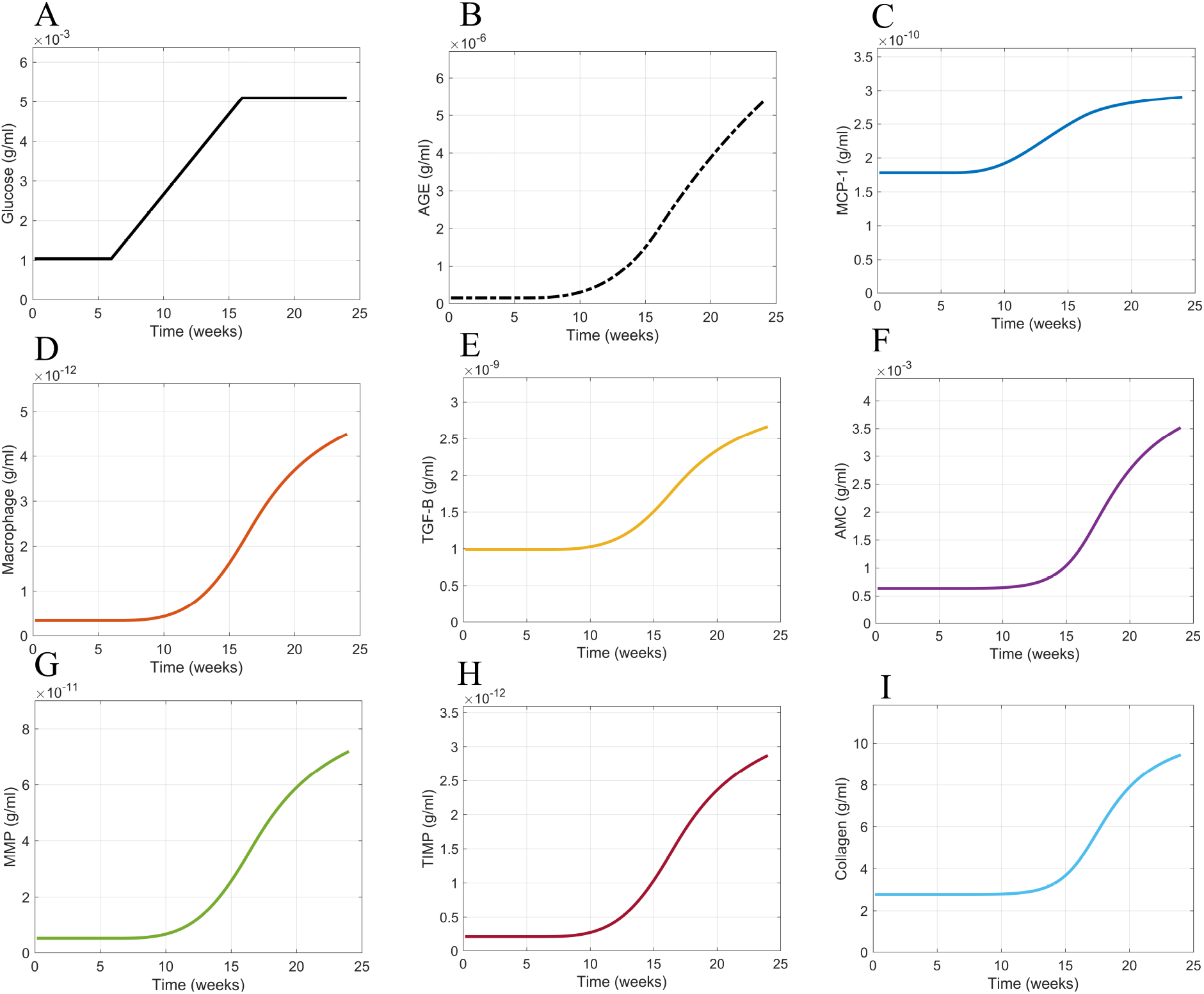
Model simulation dynamics of cells and biomolecules involved in glomerular fibrosis over 24 weeks. (A) Glucose input concentration, (B) AGE concentration, (C) MCP concentration, (D) Macrophage concentration, (E) TGF-β concentration, (F) Activated MC concentration, (G) MMP concentration, (H) TIMP concentration, and (I) collagen concentration. Abbreviations: AGE, advanced glycation end products; MCP, monocyte chemoattractant protein; TGF-B, transforming growth factor–β; Activated MC, activated mesangial cells; MMP, matrix metalloproteinase; TIMP, tissue inhibitor of metalloproteinase.

For the given conditions, these results showed that a switch in blood glucose concentration from baseline levels (0–6 weeks) to high blood glucose concentration levels (16–24 weeks) resulted in a similar switch from healthy glomerular tissue (0–16 weeks) to a fibrotic glomerular tissue (after 16 weeks) (Figure 9I).

### 4.2 Glucose control scenario

Once we had the cellular and biomolecular dynamics of glomerular fibrosis as the base case scenario, we ran different scenarios to determine what mechanistic pieces are responsible for the delayed recovery from glomerular fibrosis after achieving good blood glucose control. Good blood glucose control is defined as regulation that returns the blood glucose concentration to sustained baseline levels.

We simulated a scenario applying good blood glucose control (Figure 10A) to the model at 24 weeks via an immediate decrease in blood glucose levels from the elevated state to the normal baseline levels (Figure 10B,E). The reduction of blood glucose levels showed little effect on collagen within 30 weeks, which we defined as the short-term period (Figure 10C). Additionally, the profile of collagen was similar to that of the base case scenario (Figure 9I). For immediate blood glucose regulation, there is not an immediate decrease in glomerular fibrosis. We then simulated the model for an extended period of time (Figure 10D–G). It takes a long time (≈ 80 weeks ≈ .25 years) for glomerular fibrosis to be reversed in our *in silico* model for mice (Figure 10F). The model predicts that immediate regulation of blood glucose control does not lead to immediate amelioration of glomerular fibrosis. This agrees with the clinical observation that good blood glucose regulation via pancreas transplantation does not lead to an immediate reversal of glomerular fibrosis (Fioretto et al., 1998) but takes many months or years for the complete reversal.

**Figure 10.**
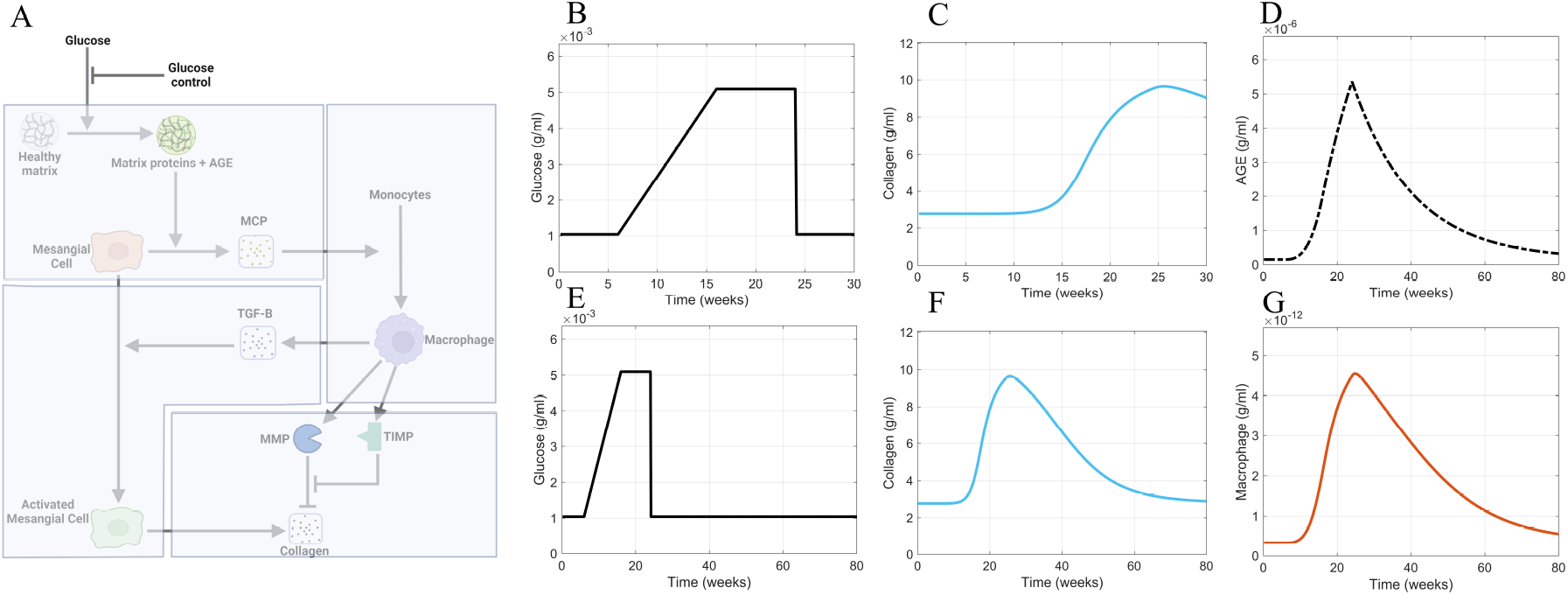
Glucose control results. (A) Non-shaded regions of the glomerular fibrosis network show changes to the model for the blood glucose control scenario (inhibition of glucose input). The effects of good blood glucose control on (B) glucose concentration and (C) collagen concentration in the short term. The effects of good blood glucose control on (D) AGE concentration, (E) glucose concentration, (F) collagen concentration, and (G) macrophage concentration in the long term. Panel A created with BioRender.com. Abbreviations: AGE, advanced glycation end products.

The continuously elevated AGE concentration caused the delayed recovery from fibrosis in the simulations. Although glucose concentration is immediately brought down to healthy-baseline levels (Figure 10B,E), the AGE concentration (Figure 10D) immediately downstream of glucose in the progression of fibrosis does not decrease to healthy baseline levels in the short term. Instead, a slow decline in the AGE concentration within the glomerulus is observed (Figure 10D). The continued elevated level of AGE causes a persistent stimulation of mesangial cells, which results in the persistent recruitment and accumulation of macrophages (Figure 10G). The accumulation of macrophages results in persistent glomerular fibrosis through the continued activation of mesangial cells, the source of the excess collagen (Figure 10F).

### 4.3 Inhibited AGE production scenario

Having determined that the continuously elevated AGE concentration is the cause of the delay in the recovery from glomerular fibrosis in our model, we sought to avert these elevated AGE levels. As our first approach, we inhibited the production of AGE (Figure 11). To implement this inhibition, the AGE equation (Equation (2)) was modified as

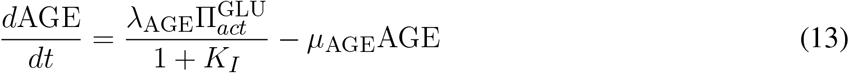

where an inhibition term 1 + *K*_*I*_ was included in the AGE formation term.

**Figure 11.**
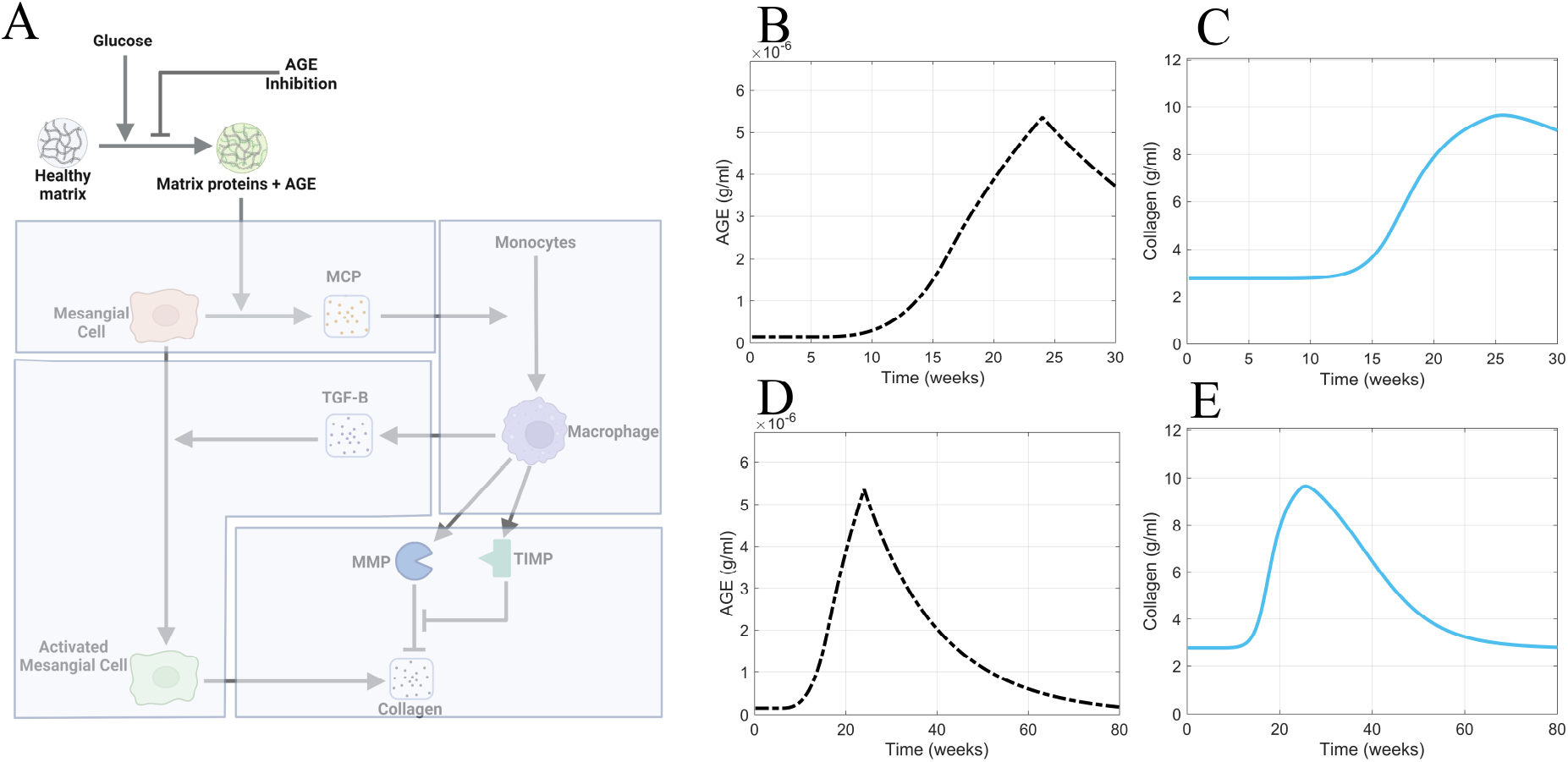
AGE inhibition is insufficient for accelerating recovery from glomerular fibrosis. (A) Non-shaded regions of the glomerular fibrosis network show changes to the model for the AGE inhibition scenario. The effects of inhibited AGE production on (B) AGE concentration and (C) collagen concentration in the short term. The effects of inhibited AGE production on (D) AGE concentration and (E) collagen concentration in the long term. Panel A created with BioRender.com. Abbreviations: AGE, advanced glycation end products.

Consistent with the other scenarios, inhibited AGE production was applied at 24 weeks (Figure 11A). Inhibited AGE production had little impact on glomerular fibrosis in the short term (Figure 11C) and took over 54 weeks for a complete reversal to occur, as seen in the long-term simulation (Figure 11E). Further, the dynamics of AGE in this scenario (Figure 11B,D) mirrored the dynamics of AGE in the glucose control scenario (Figure 10D), indicating that inhibited AGE production has the equivalent effect of applying glucose control. As such, its efficacy is limited and does not accelerate the recovery from glomerular fibrosis.

Inhibited AGE production is ineffective at immediately reversing glomerular fibrosis because glomerular fibrosis has developed at this point, and AGE is also significantly accumulated. Inhibited AGE production only prevents the further production of AGE and does not remove the already accumulated levels of AGE, thus delaying the recovery from glomerular fibrosis. Consequently, the best approach to accelerate recovery from fibrosis is to actively remove the already accumulated AGE from the system.

### 4.4 Enhanced AGE degradation scenario

To test the *in silico* hypothesis that active removal of AGE accelerates recovery from fibrosis, we simulated the model with enhanced removal of AGE from the system (Figure 12A).

**Figure 12.**
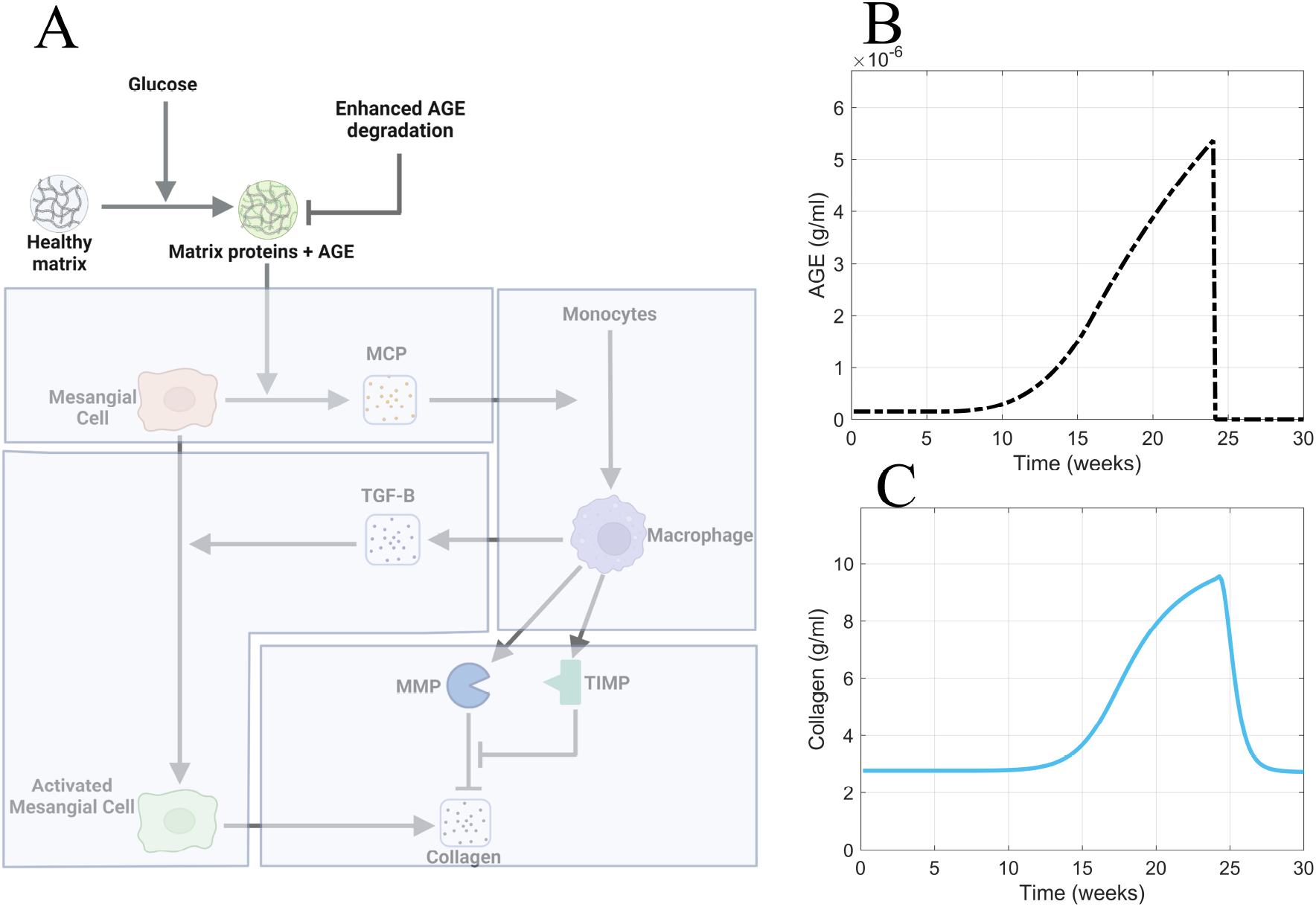
Enhanced AGE degradation is the most efficacious approach for accelerating recovery from glomerular fibrosis. (A) Non-shaded regions of the glomerular fibrosis network show changes to the model for the enhanced AGE degradation scenario. The effects of enhanced AGE degradation on (B) AGE concentration and (C) glomerular fibrosis in the short term. Panel A created with BioRender.com. Abbreviations: AGE, advanced glycation end products.

To implement the enhanced degradation of AGE, the AGE equation was modified as

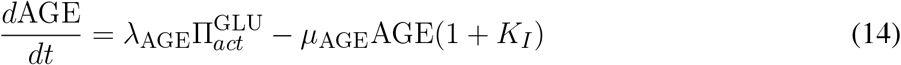

where the degradation term of AGE was multiplied by the term 1 + *K*_*I*_ to increase AGE degradation. The treatment was applied at 24 weeks, representing the enhanced degradation of AGE. In the short term, the enhanced degradation of AGE resulted in the immediate return of AGE concentration to healthy baseline levels (Figure 12B) and accelerated recovery from fibrosis (Figure 12C).

## 5 DISCUSSION

Using our model, we considered the effects of different treatment strategies for advanced glomerular fibrosis. Specifically, we asked why it takes so long to reverse the condition after glucose treatment.

Our model captured delayed recovery from advanced glomerular fibrosis when glucose control was applied (Figure 10E). This behavior was observed in patients with advanced glomerular fibrosis when glucose control via pancreatic transplant did not result in immediate recovery from fibrosis but took a long time (10 years) (Fioretto et al., 1998). The complete reversal of advanced glomerular fibrosis has not been shown in animal models, likely because the length of the studies is not long enough (Pugliese et al., 1997). The length of study for normoglycemia post-advanced glomerular fibrosis in the rat study by Pugliese et al. (1997) was a maximum of four months, during which no reversal of glomerular fibrosis was observed. However, non-advanced glomerular fibrosis has been shown to be at least partially reversible (Mauer et al., 1974, 1975; Lee et al., 1974; Steffes et al., 1980). Our model predicted that it should take around 1.25 years for a complete reversal of glomerular fibrosis to be observed (Figure 10F). Although we cannot compare this value to any data for a time to fully reverse advanced glomerular fibrosis in rodents, the value shows that it takes a long time for glomerular fibrosis to be reversed (Fioretto et al., 1998).

Our mechanistic explanation for why the reversal of advanced fibrosis takes a long time is that the slow degradation of AGE results in the persistent stimulation of mesangial cells and the continued recruitment of macrophages, which mediates the continued accumulation of collagen through the persistent activation of mesangial cells (Figure 10). The slow degradation of AGE, specifically CML, is well known. Diabetic rats that were given pancreatic transplants to achieve normoglycemia after having had diabetes for some time showed that the collagen glycated AGE content did not decrease even after four months of normoglycemia, regardless of whether glomerular fibrosis was in the early stage or advanced stage (Pugliese et al., 1997). Our model predicted a faster degradation of AGE ( 40% reduction in AGE four months post normoglycemia) than observed in these rats. Even then, we see that the reversal of glomerular fibrosis takes a long time. Our model prediction of the persistent accumulation of intermediary species, such as MCP, macrophages, TGF-β, and activated mesangial cells, even after normoglycemia is achieved, has not been studied experimentally and is still an avenue to be explored.

Having found that the slow degradation of AGE is a cause for the delay in the reversal of fibrosis in our model, we attempted to accelerate the process by inhibiting AGE formation. We found that inhibition of AGE formation did little to accelerate recovery from fibrosis because long-lived AGE was still accumulated in the glomerulus. Aminoguanidine is a molecule that inhibits the formation of AGE and has been studied as a potential therapeutic for treating glomerular fibrosis. Although the efficacy of aminoguanidine on advanced glomerular fibrosis has not been studied, its effect on the development of glomerular fibrosis in diabetic rats has been studied (Kelly et al., 2001). The study found that aminoguanidine reduces TGF-β gene expression within the glomeruli and AGE and collagen protein expression within the kidney cortex (Kelly et al., 2001). Without the complete inhibition of AGE formation, our model replicated the experimentally observed results of a reduced TGF-β expression and reduced glomerular fibrosis.

Our model predicted that the complete recovery from fibrosis with AGE formation inhibition takes 1.25 years, which means that not much recovery will be observed in a short-time frame. Clinical trials for AGE formation inhibitors, such as aminoguanidine, have had limited success (Bolton et al., 2004; Williams et al., 2007; Rabbani et al., 2009). One reason could be that once glomerular fibrosis has developed, inhibiting AGE formation is not an effective strategy to reverse glomerular fibrosis quickly. Since these clinical trials do not run for long periods, the efficacy observed could be small and not statistically significant.

The approach that we determined is the most productive at quickly reversing advanced glomerular fibrosis is the active removal of the long-lived AGE from the glomerulus. Alagebrium, an AGE crosslink breaker capable of breaking down long-lived AGE such as CML, has been studied as a potential therapeutic target for treating glomerular fibrosis. A study on diabetic rats showed that the early treatment of these rats (16 weeks into diabetes) with alagebrium for 16 weeks was able to reverse glomerular fibrosis almost to normal levels and also reduce the long-lived AGE, TGF-β, and collagen IV within the renal cortex (Forbes et al., 2003). However, the late treatment of these diabetic rats (24 weeks into diabetes) with alagebrium for 8 weeks reduced the AGE (CML) levels to baseline but did not mitigate glomerular fibrosis by a statistically significant amount, nor were TGF-β and collagen IV within the renal cortex reduced by statistically significant amounts (Forbes et al., 2003). These findings seem to disagree with our prediction that the enhanced breakdown of long-lived AGE accelerates the reversal of advanced glomerular fibrosis (Figure 12). However, our model predicted that the breakdown of AGE was followed by a rapid decrease in TGF-β, which the experimental results do not show. The absence of a reduction in TGF-β could explain why a reversal of advanced glomerular fibrosis was not observed because TGF-β, which is the key profibrotic growth factor that mediates fibrosis, was not reduced. Our model did not capture the result that the breakdown of AGE *in vivo* results in a continued accumulation of TGF-β rather than the expected reduction. The lack of decrease in TGF-β could be due to either the presence of another long-lived AGE that was not degraded by alagebrium or the breakdown of CML led to the release of bound TGF-β from the collagen matrix. Such elevation of TGF-β when a diabetic rat was treated with alagebrium has also been observed in other studies (Lassila et al., 2004), and, as such, may be an important mechanism to further explore experimentally and through computational modeling.

A phenomenon our model did not seek to replicate is the different responses to treatment between early and advanced glomerular fibrosis. Early treatment of glomerular fibrosis, such as four months after the development of diabetes, has been shown to be fully reversible within four months of normoglycemia (Pugliese et al., 1997) and late treatment of glomerular fibrosis (eight months post-induction of diabetes) has been shown to be not reversible with four months of treatment (Pugliese et al., 1997). A similar pattern has been observed when crosslink breaker alagebrium was used. The investigators observed that early treatment (four months post diabetes) reversed the glomerular fibrosis, but late treatment (6 months post diabetes) did not (Forbes et al., 2003). A potential explanation for the different responses to treatment is that the early and late stages of glomerular fibrosis occur via other mechanisms. Possibly, short-lived AGE mediates the early development of glomerular fibrosis, and long-lived AGE mediates the late development of fibrosis. Thus, early glomerular fibrosis is reversible because short-lived AGE is quickly removed from the glomerulus due to the short half-life. In contrast, late glomerular fibrosis is not quickly reversible due to the long half-lives of long-lived AGE. This hypothesized mechanism could be explored in future iterations of the model.

## 6 CONCLUSIONS

In conclusion, we developed a computational model of glomerular fibrosis in diabetes to understand the current lack of therapeutic efficacy and to propose more efficacious therapeutic approaches. Our model recapitulates the experimentally observed phenomenon that good blood glucose control does not lead to immediate recovery from glomerular fibrosis in diabetes. We determined using our model that good glucose control is not immediately efficacious due to the accumulation of AGE and slow removal from the system. We further proposed and simulated the more efficacious treatment approach of enhanced AGE degradation, which theoretically accelerates the recovery from glomerular fibrosis.

## AUTHOR CONTRIBUTIONS

HYT: Conceptualization, Data curation, Formal analysis, Investigation, Methodology, Visualization, Writing-original draft, Writing-review and editing. ANFV: Conceptualization, Formal analysis, Funding acquisition, Methodology, Project administration, Resources, Supervision, Writing-review and editing.

## Acknowledgments

We thank Dr. Stelios T. Andreadis, Dr. Rudiyanto Gunawan, and Dr. Mohammad Aminul Islam from the University at Buffalo for discussions about the model and lab members, particularly Eduardo A. Chacin Ruiz and Krutika Patidar, for their thorough feedback on the manuscript.

## DATA AVAILABILITY STATEMENT

We have provided the code and analysis files for the glomerular fibrosis model in a repository at https://github.com/ashleefv/GlomerularFibrosis (Thomas and Ford Versypt, 2023).

