## Supplementary Material for "A Mathematical Model of Glomerular Fibrosis in Diabetic Kidney Disease to Predict Therapeutic Efficacy"

### 1 DATA GATHERING

The glomerular fibrosis model was parameterized to experimental data from the literature. Here, we discuss the literature data-gathering approach.

*In vivo* data for constraining the glomerular fibrosis model was solely gathered from db/db mouse model studies. Db/db mice develop diabetes spontaneously after 6–8 weeks, as observed by their hyperglycemic condition, which arises at the ages of 4–6 weeks. These mice are used as a model for studying type 2-dependent diabetic kidney disease. They show significant mesangial matrix expansion and albuminuria, critical features of diabetic kidney disease in humans. Db/db mice are considered a good model for studying diabetic kidney disease (Hirose et al., 1982). Data from db/db mice is readily available relative to other animal models and scarce human data.

All of the *in vivo* data came from the same mouse model and not other animal models because there is significant variation in data across different animal models. In addition to variations in data across animal models, there are variations across measurement techniques and the locations of tissue samples from which data is gathered. Thus, to have consistent data, we only included data for fibrosis where the same or similar measurement techniques were used and only included data from one tissue, namely from the glomerulus of the db/db mice.

Data was gathered from the literature to connect hyperglycemia, the main stimulus for diabetes-induced glomerular fibrosis, to quantities involved in the progression of glomerular fibrosis. Time-series data for blood glucose levels were readily available. However, quantities involved in glomerular fibrosis were more sparse. The quantities involved in glomerular fibrosis for which we could find data were the time-series fold changes for glomerular collagen IV protein and TGF-β protein and cell population evolution for macrophages and activated mesangial cells. When we could not find consistent data *in vivo* from db/db mice, we also considered measurements from *in vitro* glomerular fibrosis experiments with cultured mesangial cells.

The quantities for collagen and TGF-β protein are fold change quantities, not absolute protein concentration values. This was due to the unavailability of time-series concentration data for the proteins and growth factors that are produced *in vivo* that regulate the process of glomerular fibrosis. Such data is unavailable because there are no experimental techniques to accurately quantify absolute concentration values of proteins and growth factors that are extracted from the glomerular tissue of mice. The available data is non-absolute fold change data obtained from western blot analysis and other techniques. As such, the data for collagen and TGF-β is quantified via immunohistochemical studies that use fluorescent antibodies to indicate the fold change in protein and growth factor concentration over time.

The data-gathering approach is discussed in further detail below.

### 1.1 Glucose data

Since we aimed to create a model for glomerular fibrosis in diabetes, we needed high blood glucose to stimulate the fibrosis. The idea that hyperglycemia is the main driver of glomerular fibrosis is well-established. To constrain our model, we required blood glucose concentration data in relation to the extent of glomerular fibrosis. As such, we gathered published blood glucose concentration time-series data for db/db mice from multiple sources in the literature (Figure S1) (Cohen et al., 1995, 2001, 2002; Ziyadeh et al., 2000; Koya et al., 2000; Kolavennu et al., 2008). Across the various sources, there is good consistency in the time-series blood glucose concentration within db/db mice even when different labs made the measurements. Glycemia initially increases around six weeks from baseline levels and plateaus at a high blood glucose concentration around 20 weeks (Figure S1). This data set gives quantitative dynamics for the blood glucose concentration profile within a diabetic mouse, which we used in our model as our stimulus for the progression of glomerular fibrosis (Equation (10)).

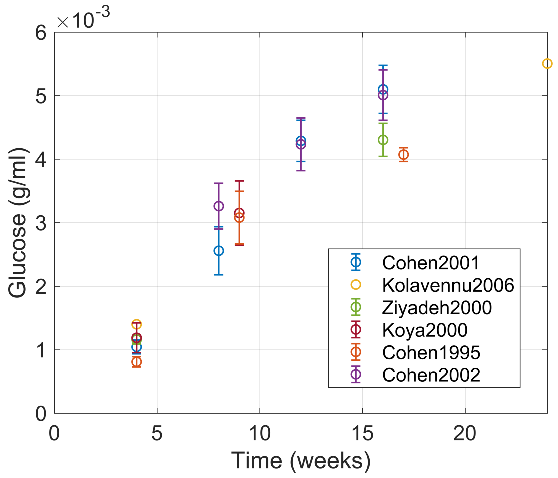


**Figure S1.** Time-series data for blood glucose concentration from db/db and control mice obtained from different sources (Cohen et al., 1995, 2001, 2002; Ziyadeh et al., 2000; Koya et al., 2000; Kolavennu et al., 2008).

### 1.2 AGE-MCP data

We gathered data from the literature (Lu et al., 2004) for dose-dependent MCP-1 response of mesangial cells to AGEs. The data were used to constrain the AGE and MCP dynamics in the disease initiation step of the fibrotic mechanism. The data were collected from an *in vitro* study of mesangial cells cultured in different concentrations of AGE to determine their MCP response (Lu et al., 2004). The data show a dose-dependent increase in the MCP expression by mesangial cells (Figure S2). *In vivo* data for AGE dynamics that were consistent with each other were not available. As such, we had to use *in vitro* data to estimate parameters in the model.

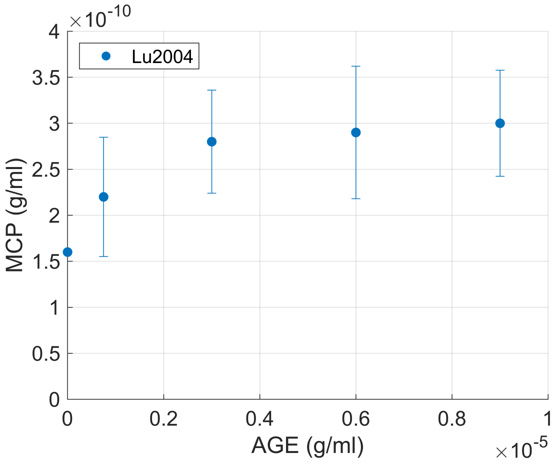


**Figure S2.** Dose-dependent MCP response of mesangial cells that were incubated in different concentrations of AGE (Lu et al., 2004). Abbreviations: AGE, advanced glycation end products; MCP, monocyte chemoattractant protein.

### 1.3 Macrophage data

We gathered data from the literature for the macrophage number fold change per glomerular cross-section in db/db mice at different time points over 20 weeks (Figure S3) (Ichinose et al., 2006; Saito et al., 2011; Hong et al., 2014; Kim et al., 2013, 2018; Choi et al., 2018; Hwang et al., 2019). The numbers of macrophages per glomerular cross-section were determined by counting the numbers of F4/80 positive cells. F4/80 is a unique marker of mice macrophages and thus allows quantifying the number of macrophages recruited to the glomerulus. The number of macrophages recruited to the glomerulus gradually increases to plateau at about 10-fold from the initial number of macrophages in healthy glomeruli (Figure S3).

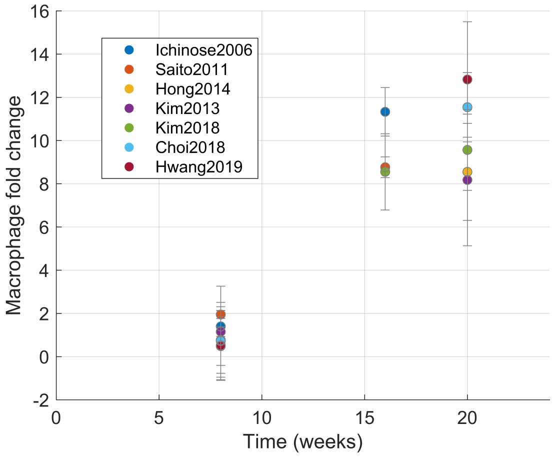


**Figure S3.** Macrophage number per glomerular cross-section in db/db mice for multiple time points in 20 weeks (Ichinose et al., 2006; Saito et al., 2011; Hong et al., 2014; Kim et al., 2013, 2018; Choi et al., 2018; Hwang et al., 2019).

While gathering the data from the literature, we only used data from experimental studies that used F4/80 as a marker and not other markers such as CD68. CD68 is nonspecific between monocytes and macrophages and is also a standard marker for tumor-associated macrophages (Harris et al., 2012). Additionally, the data from CD68 studies were inconsistent with those of the F4/80 (Seo et al., 2015). We also excluded experimental studies that did not inspect a large enough sample of glomeruli per mouse (minimum of 20 glomeruli per mouse) because small samples also led to inconsistent data points (Terami et al., 2014).

### 1.4 TGF-β1 protein area fold change data

We gathered data from the literature for TGF-β dynamics. TGF-β1 protein area fold changes per glomerular cross-section were measured by immunostaining the glomerular tissues with anti-TGF-β1 antibodies, and then the area fold changes were scored semi-quantitatively (Figure S4) (Park et al., 2016; Hong et al., 2014; Kim et al., 2018; Choi et al., 2018). The TGF-β protein area fold change was the only type of data available that quantified the amount of TGF-β protein expressed during glomerular fibrosis.

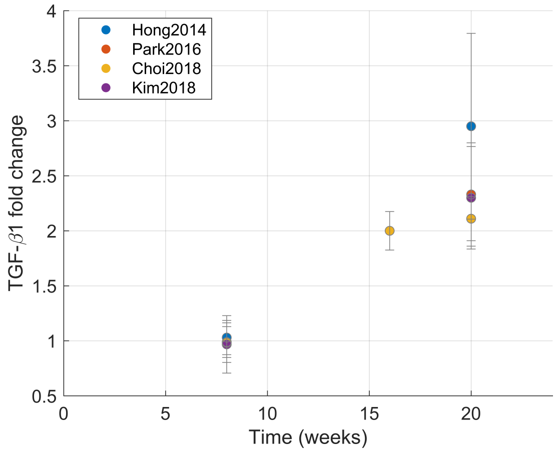


**Figure S4.** TGF-β1 protein score fold change data for db/db mice per glomerular cross-section over 20 weeks. Data was obtained from multiple sources for TGF-β dynamics in db/db mice glomeruli (Hong et al., 2014; Park et al., 2016; Choi et al., 2018; Kim et al., 2018). Abbreviations: TGF-B1, transforming growth factor–β1.

### 1.5 Activated mesangial cell data

We gathered $\alpha$-smooth muscle actin ($\alpha$-sma) expression per glomerular cross-section over 20 weeks within the db/db mice, and $\alpha$-sma is an indicator of activation of mesangial cells. It is used as a proxy for the fold change of the population of activated mesangial cells (Figure S5A) (Wang et al., 2016). We also gathered dose-response data for $\alpha$-sma expression in mesangial cells incubated in varying concentrations of TGF-β (Figure S5B) (Fu et al., 2013).

### 1.6 Collagen IV protein area fold change data

Collagen IV protein area fold changes in the glomerulus were determined by immunostaining using anti-collagen IV antibodies and different morphometric techniques (Figure S6) (Ichinose et al., 2006; Kosugi et al., 2010; Kim et al., 2013; Chen et al., 2014; Hong et al., 2014; Park et al., 2016; Choi et al., 2018; Kim et al.,

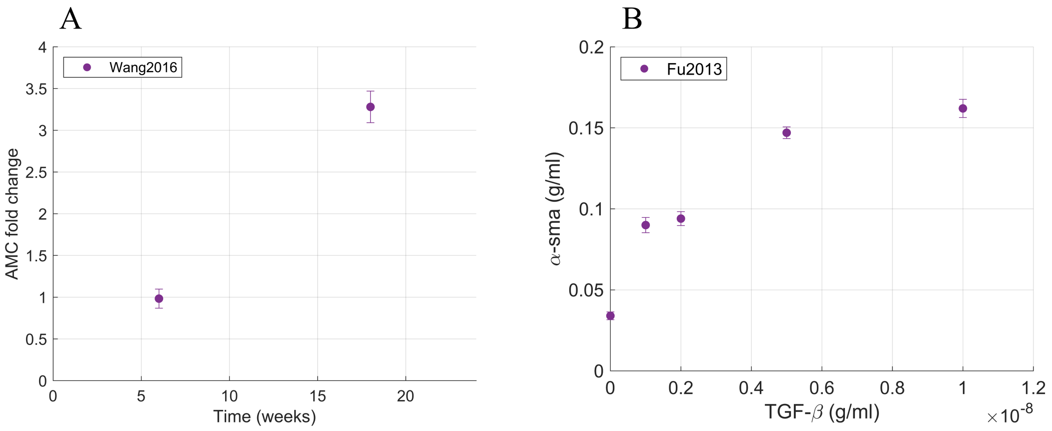


**Figure S5.** Data relevant to activated mesangial cells. (A) Activated mesangial cell population fold change per glomerular cross-section for db/db mice over 20 weeks (Wang et al., 2016). (B) Dose-dependent $\alpha$-sma expression in mesangial cells incubated in different concentrations of TGF-β (Fu et al., 2013). Abbreviations: $\alpha$-sma, $\alpha$-smooth muscle actin.

2018; Hwang et al., 2019). The collagen IV protein area was quantified using semi-quantitative measurement techniques (blinded scoring) and quantitative morphometric methods. Data from both measurement techniques were combined because they were consistent relative to each other, which enabled the filling of gaps in the separate datasets. Area fold change data was collected because it is the only type of collagen IV glomerular protein data available in the literature. There are quantitative collagen IV mRNA measurements done by isolating glomeruli and quantifying mRNA amounts using reverse transcription polymerase chain reaction (RT-PCR) (Wang et al., 2016; Nagai et al., 2019; Park et al., 2014; Hsu et al., 2021), but these data cannot be used to represent collagen IV protein dynamics because mRNA concentration does not necessarily translate directly to protein concentration. The different glomeruli isolation techniques used to isolate collagen IV mRNA from only glomeruli, such as laser capture microdissection (Hsu et al., 2021), magnetic bead method (Nagai et al., 2019; Li et al., 2020), and sieving method (Park et al., 2014), could be very useful techniques in the future to isolate proteins from the glomerulus to enable the quantification of absolute concentration values.

### 2 SENSITIVITY ANALYSIS

The sensitivity of peak collagen accumulation to the parameters (Figure S7A) indicates the parameters that influence the extent of fibrosis most. The results show parameters associated with MCP dynamics, activated mesangial cell dynamics, and TGF-β dynamics have the strongest influence on the amount of collagen accumulated (Figure S7A). Out of all those parameters, the Hill function parameters $n_{MCP}$ and $K_{MCP}$ for the recruitment of macrophages and the degradation rate of MCP $\mu_{MCP}$ are the most influential negative regulators on the extent of collagen accumulation. Then, the mesangial cell population $MC$, the sources of MCP $S_{MCP}$ and $\lambda_{MCP}$, and the sources of TGF-β and activated mesangial cells $\lambda_{TGF}$ and $\lambda_{AMC}$ are the most influential positive regulators of collagen accumulation. The significant influence of MCP dynamics on collagen accumulation could be due to the MCP-mediated macrophage recruitment being the only mechanism within the model for fibrosis

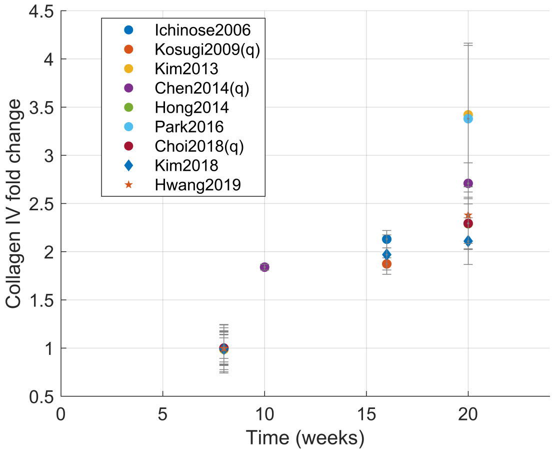


**Figure S6.** Collagen IV protein area fold change data from db/db mice over 20 weeks. Data was obtained from multiple sources (Ichinose et al., 2006; Kosugi et al., 2010; Kim et al., 2013; Chen et al., 2014; Hong et al., 2014; Park et al., 2016; Choi et al., 2018; Kim et al., 2018; Hwang et al., 2019).

to progress. Adding other mechanisms could improve model predictions, such that fibrosis is not highly dependent on MCP dynamics.

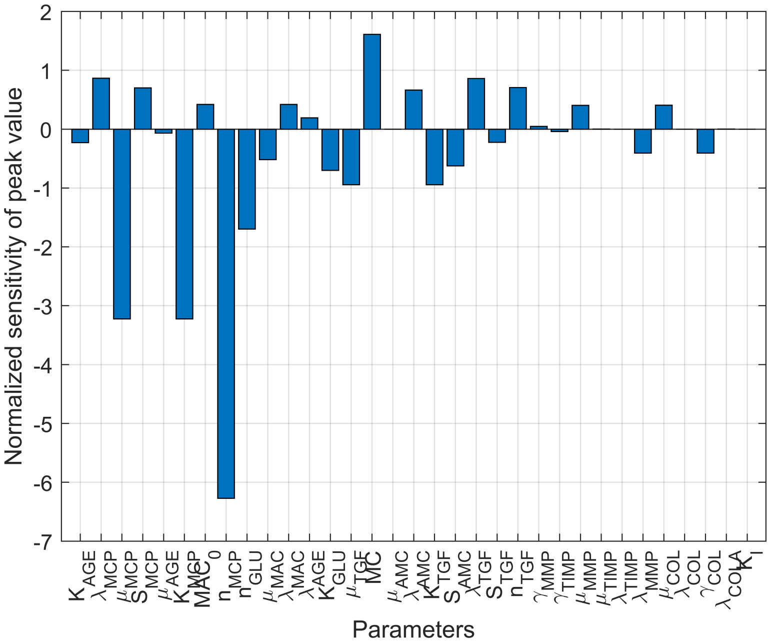


**Figure S7.** Local sensitivity analysis results. Normalized sensitivity of peak value of collagen concentration. Abbreviations: AGE, advanced glycation end products; MCP, monocyte chemoattractant protein; MC, mesangial cells; MAC, macrophages; TGF, transforming growth factor–β; AMC, activated mesangial cells; MMP, matrix metalloproteinase; TIMP, tissue inhibitor of metalloproteinase; COL, collagen. Other notation defined in Table 1.

*Proceedings of the National Academy of Sciences USA* 97, 8015–8020. doi:10.1073/pnas.120055097
